# Candidate genes associated with neurological manifestations of COVID-19: Meta-analysis using multiple computational approaches

**DOI:** 10.1101/2022.04.10.487761

**Authors:** Suvojit Hazra, Alok Ghosh Chaudhuri, Basant K. Tiwary, Nilkanta Chakrabarti

## Abstract

COVID-19 develops certain neurological symptoms, the molecular pathophysiology of which is obscure. In the present study, two networks were constructed and their hub-bottleneck and driver nodes were evaluated to consider them as ‘target genes’ followed by identifying ‘candidate genes’ and their associations with neurological phenotypes of COVID-19. A tripartite network was first constructed using literature-based neurological symptoms of COVID-19 as input. The target genes evaluated therefrom were then used as query genes to identify the co-expressed genes from the RNA-sequence data of the frontal cortex of COVID-19 patients using pair-wise mutual information to genes. A ‘combined gene network’ (CGN) was constructed with 189 genes selected from TN and 225 genes co-expressed in COVID-19. Total 44 ‘target genes’ evaluated from both networks and their connecting genes in respective networks were analyzed functionally by measuring pair-wise ‘semantic similarity scores’ (SSS) and finding Enrichr annotation terms against a set of genes. A new integrated ‘weighted harmonic mean score’ was formulated using SSS and STRING-based ‘combined score’ to select 21 gene-pairs among ‘target genes’ that provided 21 ‘candidate genes’ with their properties as ‘indispensable driver nodes’ of CGN. Finally, six pairs providing seven prevalent candidate genes *(ADAM10, ADAM17, AKT1, CTNNB1, ESR1, PIK3CA, FGFR1)* exhibited direct linkage with the neurological phenotypes under tumour/cancer, cellular signalling, neurodegeneration and neurodevelopmental diseases. The other phenotypes under behaviour/cognitive and motor dysfunctions showed indirect associations with the former genes through other candidate genes. The pathophysiology of ‘prevalent candidate genes’ has been discussed for better interpretation of neurological manifestation in COVID-19.

## Introduction

The ‘coronavirus disease 2019’ (COVID-19) patients present common symptomatic features of dry cough, dyspnea, fever, fatigue and myalgia^1,2^ followed by the acute respiratory distress syndrome (ARDS) in an advanced stage^3^. The COVID-19 is caused by severe acute respiratory syndrome coronavirus 2 (SARS-CoV-2), which belongs to the genera of β-coronavirus having positive single-stranded RNA as its genome^3–6^. The SARS-CoV-2 can infect a wide variety of human tissue cells, leading to a greater viral load resulting in a multiorgan failure. The pathophysiological action of the virus always begins with the binding of spike proteins onto the angiotensin-converting enzyme 2 (ACE2) receptor proteins in the host cell membranes and expresses several phenotypic manifestations in human^7,8^.

The human ACE2 receptors are constitutively expressed in neurons, pericytes and endothelial cells of the olfactory bulb and certain neuroglial cells like astrocytes and oligodendrocytes. Neuromodulator regions of dopaminergic, serotoninergic, histaminergic and norepinephrinergic nuclei and several specific brain regions including substantia nigra and piriform cortex, are also rich in ACE2 receptors. Relatively, a greater expression of ACE2 receptors is evident in the brain stem, central glial substance and choroid plexuses of lateral ventricles of the brain9. The ACE2 levels were found to be very low in the human prefrontal cortex and hippocampus^9^. COVID-19 is reported to associate with lesions in the hippocampus^10^ and frontal cortex^11^ and, the structural changes in the brain stem, medulla and pons^12^. The SARS-CoV-2 RNA has also been detected in the medulla and cerebellum through an RT-PCR study that represented contamination in the leptomeninges and Virchow-Robin spaces (perivascular space)^13^.

The SARS-CoV-2 can enter the brain through three possible pathways via (i) the inflammatory supporting cells of the olfactory mucosa^13^, (ii) the endothelial cells of the cerebral blood vessels^14,15^ and (iii) the nerve terminals of the vagi in the respiratory^14,15^ and gastrointestinal tracts. SARS-CoV-2 was found to be present in the cerebrospinal fluid (CSF) of patients suggesting the predominance of immunological damages over the viral replication in neurons^16^. Among the three pathways, the first one appears to be the most important, as a majority of the COVID-19 case reports suggest anosmia and ageusia^17^ as the non-specific symptoms^18,19^. The clinical reports and the neuroimaging studies suggest that the cytokine storm and oxidative stress along with the reduction of GSH levels are two key mechanisms that can produce neurodegenerations in certain areas of the human brain^12,20^. Certain review reports^21–24^ speculate that COVID-19 related symptoms can together act as direct or indirect mediators of various neurodegenerative diseases including dementia, Alzheimer’s Disease, and Parkinson’s Disease, although the exact mechanism is still in debate.

The systematic review and meta-analysis evidences^25,26^ including the latest retrospective cohort study^27^ on 2,36,379 COVID-19 survivors indicate that 33.62% of patients have neurological or psychiatric problems. A literature survey indicates that cognitive alterations (delirium with a combination of acute disturbances in attention, awareness and cognition), motor dysfunctions (dizziness, cerebellar ataxia, dysautonomia, seizure and epileptogenesis), Cerebro-vascular changes (cerebral ischemia and infarct, stroke, focal ischemic necrosis, oedema, cerebral and subarachnoid haemorrhage, subdural hematoma), Cerebro-structural changes (astrocytosis, microgliosis, meningitis/encephalitis, Encephalopathy, necrotizing hemorrhagic encephalopathy, multifocal lesions in both cerebral hemispheres, leptomeningeal enhancement, swelling in midbrain region, spinal cord myelitis are found in COVID-19 patients^28–42^. On the other hand, symptoms like myopathy^43^ and muscle injury^44^, neuropathy and polyradiculopathy including Guillain-Barré syndrome^45^, may occur in the peripheral nervous system.

In the present scenario, there is one bioinformatics-driven systems-level study using bipartite models of disease-gene, disease-disease, miRNA-gene, drug-protein interactions, which reveals that a variety of neurological symptoms including dementia, ataxia, encephalopathy and stroke along with their associated genes lined with multiple cellular functional pathways, can be therapeutically targeted by repurposed drugs or chemical compounds^46^. Additionally, a tripartite network modelling has been reported encompassing endocrine-disrupting chemicals (EDC), targeting proteins and diseases as the three types of nodes that decipher putative links between EDCs, COVID-19 severity and association to other diseases^47^. A systems-level modelling study^48^ has been conducted for the construction of a tripartite network of symptom-disease-gene to unravel the interplay between phenotype and genotype during disease conditions that are not limited to nervous system manifestation. Recent network-based findings of hub-bottleneck nodes for drug repurposing study report the involvements of several molecules associated with immunological systems (viz. cytokines e.g., TNFα, IL-1β,-6,-10 and chemokines e.g., CXCL8 and CCL2), growth factor function (e.g., VEGFA), cell-to-cell interaction (e.g., ICAM1), and signal transduction pathway (e.g., AKT1) with the neurological complication in COVID-19^49^.

In the present study, a novel approach has been introduced, for the first time to the best of our knowledge, to find a model of predictive candidate genes and their associations with neurological phenotypes of COVID-19. Initially, a tripartite network (TN) has been constructed using literature evidenced neurological symptoms of COVID-19 as input whereby integrated weightage of symptoms and diseases are implied to get the most robust predictive genes for TN. Secondly, the predictive target genes evaluated from TN have been considered as co-regulated in tissue and used as query genes to identify a set of co-expressed genes (CG) from RNA-sequence data of the frontal cortex of COVID-19 patients using pair-wise mutual information (transcriptional gene-gene interaction from expression levels) to genes. The ‘combined gene network’ (CGN) has been constructed using genes selected in TN and co-expressed genes evaluated from RNA-sequence data of COVID-19. Both networks were analyzed topologically and functionally to get ‘candidate genes’ and their connections with functional annotations to find the putative molecular pathophysiology in the brain associated with COVID-19.

## Materials and methods

The systematic methodology with inclusion and exclusion criteria applied in the present meta-analysis documented with the flow diagram following Preferred Reporting Items for Systematic Reviews and Meta-Analyses (PRISMA)^50^ guideline (Fig. 1).

**Fig. 1.**
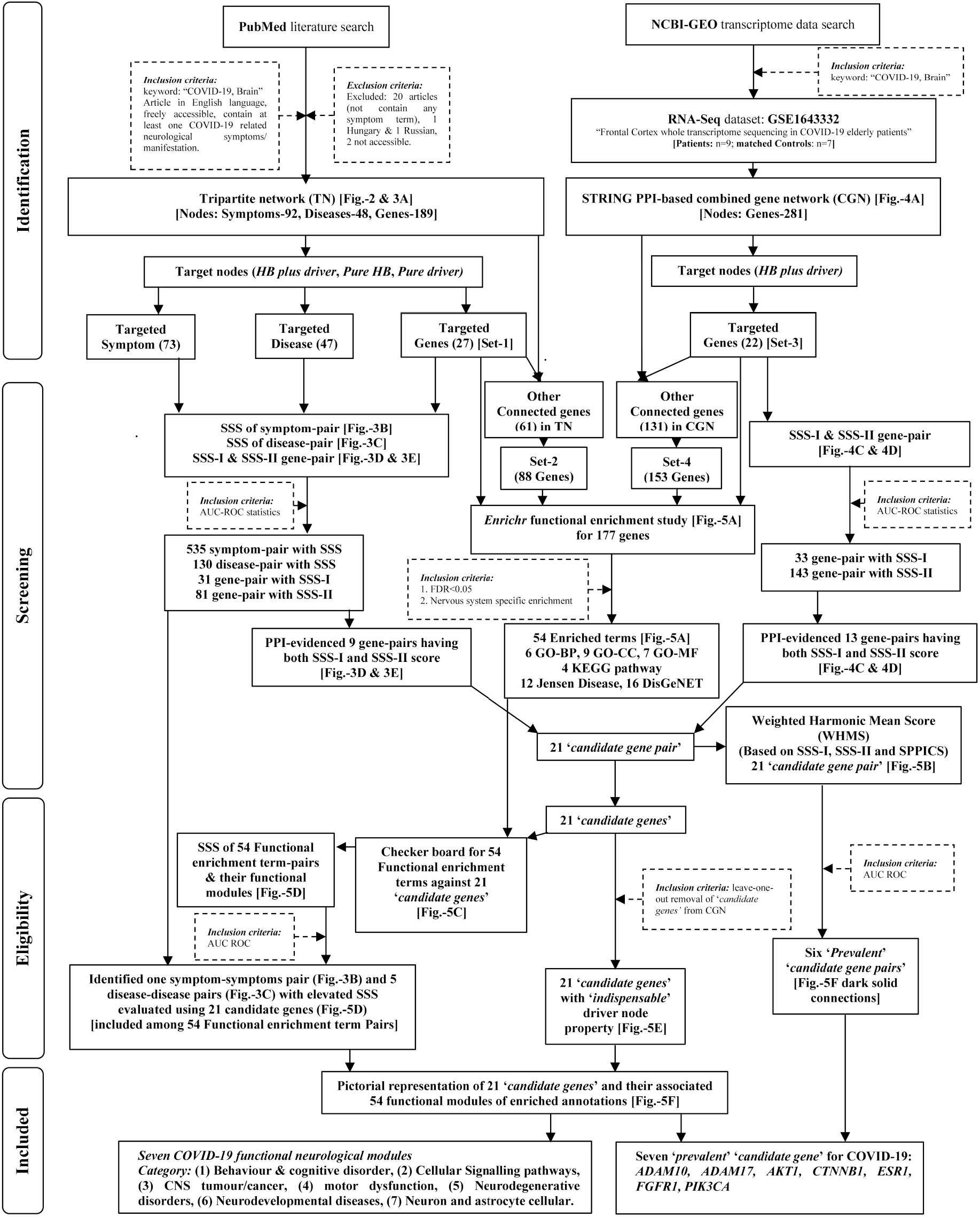
PRISMA Flowchart of the systematic and stringent methodology applied and the results found in a systems level analysis to identify *‘candidate genes’* and associated functional modules (symptoms/diseases) related to neurological manifestations of COVID-19.

### 1. *In silico* modeling of Tripartite Network (TN) for COVID-19

The TN of symptom-disease-gene (nodes) was constructed using connections (edges) developed by mathematical and statistical formulations. The stepwise approach for construction of TN is presented in the pictorial diagram (Fig. 2). The terms or keywords of neurological symptoms (represented by nodes) related to COVID-19 were found by systematic PubMed bibliographic literature search. Initially, the symptom-based disease and gene bipartite (symptom-disease and symptom-gene) networks were constructed by generating a corpus of data using Human Phenotype Ontology (HPO)^51^, a comprehensive digital vocabulary database of disease associated phenotype information including genomic interpretation of diseases.

**Fig. 2.**
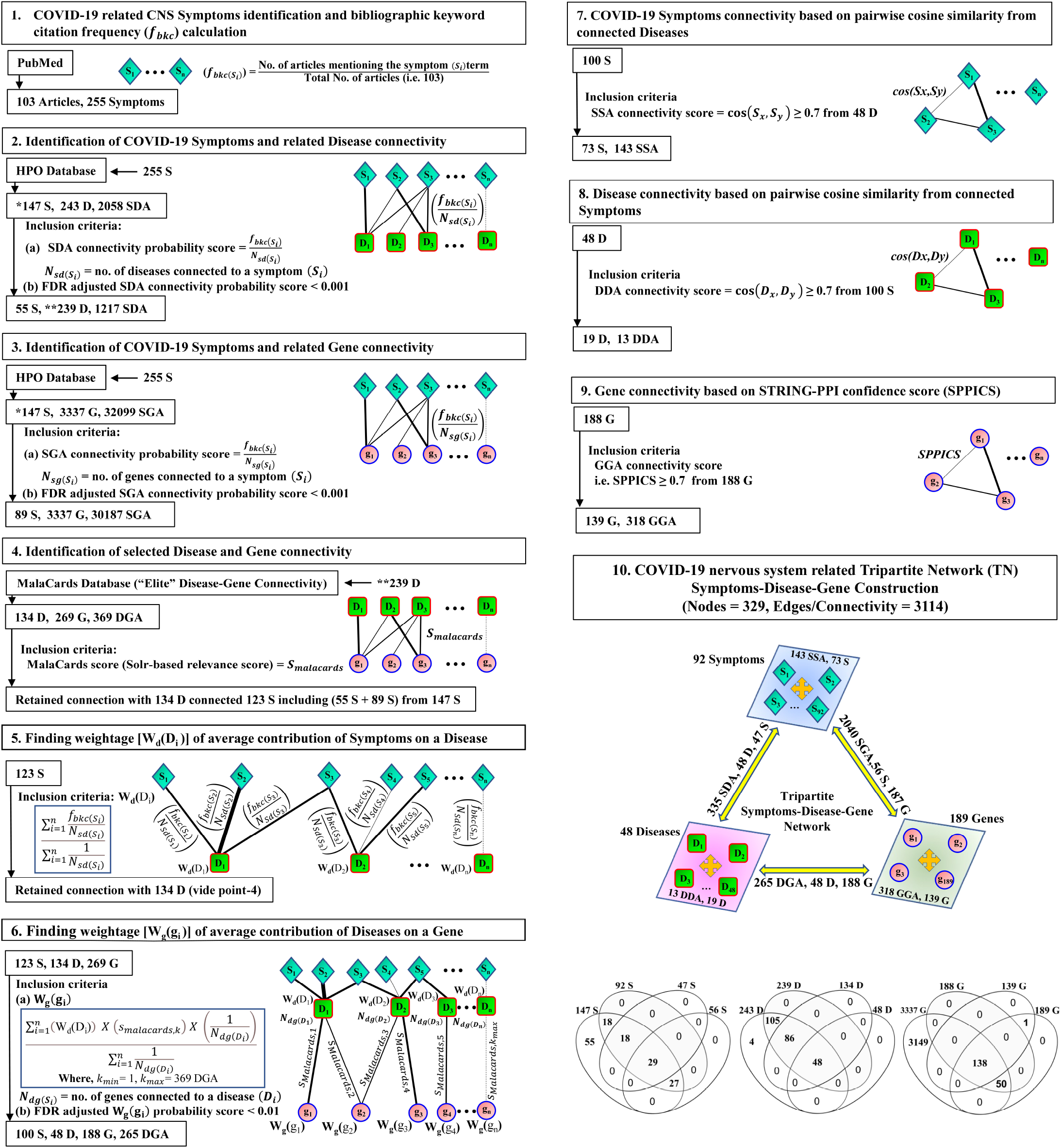
Study design for the construction of the TN for COVID-19: Stepwise presentation of construction of TN having symptoms, diseases and genes as nodes and their interactions as edges including inter-interactions (viz., symptom-disease, symptom-gene, disease-gene) in Point-1 to point-6 and intra-interactions (viz. symptom-symptom, disease-disease, gene-gene) in Point-7 to point-9 of description. Point-1: Extraction of symptoms as terms associated with neurological disorders in COVID-19 from PubMed bibliographic literature database and assigning a metric viz. bibliographic occurrence frequency (***f**_bkc_*) to each term of symptom. Point-2 and Point-3: Extraction of the interactions between symptom to disease (point-2) and symptom to gene (point-3) from HPO database using the symptom terms as query and calculating their respective connectivity probability score followed by selection of best fitted connections using the statistical FDR (p<0.001) analysis. Point-4: Extraction of the ‘Elite’ disease-gene connections with respective Solr-based relevance score (S_malacards_) from Malacards database using symptom-associated disease terms (found in point-2) as query respect to symptom-associated gene terms (found in point-3). Point-5 and point-6: Implication of symptoms on connections of disease-genes by computing ‘weightage of average contribution’ of symptoms on diseases (W_d_(D_i_)) and that of diseases on genes (W_g_(g_i_)) to find the Elite’ Disease-gene connection considering co-occurrence of at least connections of one disease to one symptom and one gene terms in the network. The FDR adjusted p<0.01 was used to filter weighted-based disease-gene connections. Point-7 to Point-9: Finding intra-connectivity of nodes using cosine semantic similarity scores ≥ 0.7 for retaining symptom-symptom (point-7) and disease-disease (Point-8) pairs and, STRING confidence score ≥ 0.7 for retaining gene-gene pairs. These symptoms, diseases and genes are selected mathematically and statistically as described in point-6. Point-10: Integration of all inter- and intra-connectivity of symptoms and their allied diseases and genes to construct the COVID-19 associated symptom-disease-gene tripartite network (TN). The representative of Venn diagrams indicates the step-wise changes in number of nodes for selection of elite symptoms, diseases and genes for construction of TN.

The bipartite networks were constructed providing weightage to each symptom i.e., *‘bibliographic keyword citation frequency’* (f_bkc(Si)_) as the count of appearance of a keyword/term in an article set^52^ and assigning its connectivity (N_sd(Si)_ or N_sg(Si)_) probability score to diseases (f_bkc(Si)_/N_sd(Si)_) and genes (f_bkc(Si)_/N_sg(Si)_) considering co-occurrence of at least one disease/gene connection to one symptom term following the principle of frequency of the co-occurrence of root node in the directed acyclic graph^53^. The false discovery rate (FDR)-adjusted p<0.001 was used to filter connections in both symptom-disease and symptom-gene bipartite network.

The new approach was implemented to find disease–gene interactions using selected diseases and genes found in bipartite networks. The Malacards^54^, a comprehensive integrated database of human maladies and their annotations including genetic entities, was mined to find the Elite disease-gene interactions with scores (Sorl’s relevance score) as the strength of their interactions. In addition, the integrated symptom-based weightage (W_d_(D_i_)) of diseases was generated (point-5 of Fig. 2) which were multiplied by respective Malacards-scores to get disease-gene ‘integrated connection scores’ followed by calculating their average to get weightage of a gene (W_g_(g_i_)) considering co-occurrence of at least connections of one disease to one symptom and one gene term in the network (point-6 of Fig. 2). The weightage (W_d_(D_i_)) of a disease was calculated by dividing the average of ‘connection probability score’ (f_bkc(si)_/N_sd(si)_) of symptoms to a disease by the summation of the frequency of connections (point-5 of Fig. 2). The FDR-adjusted p<0.01 was used to filter disease-gene connections and a tripartite network (TN) was formed. The intra-connectivity of nodes within tripartite network was further filtered by using cosine semantic similarity scores ≥ 0.70 for retaining symptom-symptom and disease-disease pairs. The pairwise cosine semantic similarity between symptoms were calculated^55^ as considering each symptom node as vector of connected diseases and vice versa for disease-disease cosine similarity calculation by considering each disease node as a vector of connected symptoms. The gene-gene intra-connectivity were filtered by using STRING^56^ confidence score ≥ 0.70 to get the final TN. The topology analysis of the TN was done in Cytoscape software^57^ using the CentiScaPe module^58^ to find ‘target nodes’ (hub-bottleneck and driver nodes) for symptoms, diseases and genes.

### 2. Finding of the co-expressed genes of COVID-19

The important nodes (Hub-bottlenecks and driver nodes) evaluated from TN were considered as co-regulated in tissue and used as query genes to identify a set of co-expressed genes from RNA-Seq data^59^ (NCBI-GEO accession ID: GSE164332) of brain frontal cortex of COVID-19 patients (n=9) with healthy matched controls (n=7), using *geneRecommender* algorithm^60,61^ in R software and environment. In this algorithm, the count data of the RNA-Seq samples of the input dataset for gene co-expression study was first normalized, then cross-validation was performed by leave-one-out method and genes were finally ranked based upon Spearman correlation with query genes using Z-score. The co-expressed genes were selected using *minet* package^62^ in R-language based on algorithm for ARACNe (Algorithm for the Reconstruction of Accurate Cellular Networks)^63^ assigning weights of (a) pair-wise mutual information (transcriptional genegene interaction from expression levels) to genes as nodes and (b) empirical probability (entropy estimators) to its edge with given threshold value for refining node-pairs.

### 3. In silico modeling of STRING-PPI network using selected co-expressed genes and genes of TN

A combined set of 225 genes collected from *minet* output (co-expressed genes) and all 189 genes of TN, were incorporated as a query in STRING database^56^ using ‘STRING-PPI combined score’ (SPPICS) 0.60 as threshold, to construct a PPI-based ‘combined gene network’ (CGN). The study was further extended to construct networks using three other different ‘SPPICS’ viz. 0.70, 0.80 and 0.90 as thresholds in respective cases. The CGNs were incorporated separately into the Cytoscape software^57^ to find HB and driver nodes for each network using the CentiScaPe module ^58^ The HB and driver nodes were cumulated from four CGNs termed as second set of *‘target genes’* evaluated from CGN.

### 4. Topology analysis of networks to find ‘hub’, ‘bottleneck’ and ‘driver’ nodes

The centrality measurements of networks were analyzed using the CentiScaPe module^58^ in Cytoscape software^57^ to find *‘hub’* (high degree) and *‘bottleneck’* (high-betweenness/shortest-path) nodes that having higher scores than cut-off values i.e., respective average node degree and average node betweenness scores (for TN: 18.93 and 465.16; for CGN: (i) SPPICS>0.60: 5.64 and 795.01, (ii) SPPICS>0.70: 4.43 and 644.21, (iii) SPPICS>0.80: 3.56 and 531.32, (iv) SPPICS>0.90: 3.17 and 406.16). The nodes which satisfied both hub and bottlenecks *(HB:* Hub-bottlenecks) properties were selected as *‘date-hubs’* considering that those nodes provided higher level of intermodular connector to co-ordinate various functional complex modules in a complex biological network^64–67^.

The controllability measurements of two networks were analyzed by finding *‘driver nodes’* using the Minimum Driver node Set (MDS) algorithm from the CytoCtrlAnalyser^68^ module of Cytoscape software. The driver nodes are the minimum number of nodes in a network which control all nodes by receiving the input signals and provide the controllability measurements to analyze the temporal (dynamic) properties of a complex network^69,70^. Driver nodes are classified into three categories based on the impact of the nodes on controlling main network viz. (a) *‘indispensable’* i.e., positive control factor, the removal of which increases the total driver nodes in main network, (b) *‘dispensable’* i.e., negative control factor, the removal of which decreases the total driver nodes in main network and (c) *‘neutral’* control factor the removal of which does not change total driver nodes in main network^69,71^. In the present study, the finally selected ‘candidate genes’ were drawn from CGN (combined gene network) by leave-one-out method to cross-validate the control property (*‘indispensable’/‘dispensable’/‘neutral’*) of the nodes on the CGN by measuring the % changes of driver nodes (total and pure driver) after removal of the specific *‘candidate genes’.* The pure driver nodes were designated as driver nodes without having hub-bottleneck properties.

The nodes having high centrality (hub with bottleneck properties) as well as controllability (driver property) values were supposed to be the disease candidates^69,72^ and therefore termed those selected nodes as *‘target symptoms’, ‘target disease’* and *‘target genes’* (TG-TN) of COVID-19 evaluated from TN. In parallel, the *‘target genes’* (TG-CGN) of COVID-19 were also evaluated from CGN.

### 5. Gene ontology-based pair-wise semantic similarity measurement of *‘target nodes’*

The functional associations among *‘target nodes’* were analyzed by semantic comparison of Gene Ontology (GO) terms^73,74^ quantitatively through computing similarities between gene-pairs and also clustering gene-pairs into known pathways. In this regard, the Wang’s GO-BP (biological process) ‘semantic similarity score’ (SSS) had been measured using *GOSemSim* package^75^ in R-language using best-match average (BMA) combination strategy to get the results closer to human expectations^76,77^. The SSS measurements were performed in two functions^75^ namely, (a) ‘*mgeneSim*’ and (b) ‘*mclusterSim*’ that estimated pairwise semantic similarities for a list of gene and ‘gene-clusters’, respectively. A ‘gene-cluster’ represented the modular nature of connected gene-set that resorted the annotation similarity network of the gene-set to prioritize the connected genes against a ‘*target node*’. Therefore, the SSS measured in two ways (a) *‘direct association’,* designated as SSS-I, by the comparative assessment of associated GO-BP terms of each of two genes and (b) ‘indirect association’, designated as SSS-II, to represent the summated contribution of comparative assessment of associated GO-BP terms of gene-clusters against targeted gene-pairs.

Notably, SSS-I and SSS-II were measured for ‘target genes’ evaluated from both TN and CGN. The pair-wise SSS (SSS-II) for *‘target diseases’* was measured based on their connecting ‘gene-clusters’ in the TN using *mclusterSim* function in *GOSemSim* package^75^ following the top-down disease module approach^48^ whereby the inference of disease-disease association was drawn based on underlying molecular mechanism. The similar approach was used for finding pair-wise SSS for *‘target symptoms’* found in the TN. Further, the classifier statistics ROC-AUC was introduced using *pROC* package^78^ in R software and environment to filter out spurious pair-wise SSSs for symptom-, disease- and gene considering the accuracy classification as excellent (0.9 < AUC < 1.0), good (0.8 < AUC < 0.9) and weak (AUC <0.8). The optimal threshold for ROC based on optimum F1 score and maximum accuracy was considered as cut-off value for selection of gene-pairs among ‘target nodes’ for finding ‘candidate nodes’ for symptoms, diseases and genes.

### 6. Functional enrichment analysis of the sets of *‘target genes’* and their connected genes

The functional annotations were found using four different gene-sets viz. (i) Set-1: *‘target genes’* evaluated from TN, (ii) Set-2: *‘target genes’* and their connected genes in TN, (iii) Set-3: *‘target genes’* evaluated from CGN and (iv) Set-4: *‘target genes’* and their connected genes from CGN. The GO terms (BP, CC, MF), KEGG pathway, disease modules (DisGeNET, Jensen Disease) analyses were performed using the *Enrichr* platform^79, 80^, an intuitive web-tool for gene over-representation study with an inclusive functional annotation set, by considering each gene-set as query. FDR-adjusted p-value less than 0.05 related to nervous system were considered as significantly enriched term. In the present study, the Set-2 and Set-4 gene-sets were pondered to double-check the over-represented functional annotation of respective Set-1 and Set-3 gene-sets. The pairwise SSS-II scores for statistically confident enriched terms were calculated using *mclusterSim* function in *GOSemSim* package^75^ as previously described methods. The functional annotation terms for *‘candidate genes’* were manually curated from integrated Enrichr results and were further analyzed for the pairwise SSS-II scores to find statistically confident enriched terms using *mclusterSim* function in *GOSemSim* package as previously described methods. The classifier statistics ROC-AUC was used to find accurate classification based on AUC values and select functionally associated Enrichr terms based on the optimal threshold for ROC as cut-off of SSS-II scores.

### 7. Formulation of integrated ‘weighted harmonic mean score’ (WHMS)

The integrated ‘weighted harmonic mean score’ (WHMS) were evaluated using harmonic mean of weightage scores for gene-pairs among *‘target genes’* which appeared to fulfil the criteria of having (a) three individual scores of SPPICS, SSS-I, SSS-II and (b) at least one score with value above respective threshold (cut-off) level. The *‘accuracy values of ROC* of SPPICS (W_SPPICS_), SSS-I (W_SSS-I_), SSS-II (W_SSS-II_) were applied as weightages for respective cases following the principle reported^81^ previously. The formula for integrated WHMS used in the study is given below.

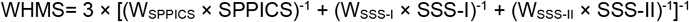

The classifier statistics ROC-AUC and the optimal threshold for ROC as cut-off of WHMS were utilized to obtain the prevalent *‘candidate gene-pairs’* considering them as putative disease associated genes.

## Results

The results found in the present meta-analysis of the rapport of brain disease related phenotypes and molecular machineries involving genes/proteins were systematically documented in the flow diagram with the findings of prevalent *‘candidate genes’* and their links with neurological functional modules in COVID-19 (Fig. 1).

### 1. Construction of TN using symptom, diseases and genes for COVID-19

Among 255 symptoms of COVID-19 patients obtained from 103 selected articles under PubMed search, the 147 symptoms linked with 243 diseases and 3337 genes having connections of 2058 symptom-disease and 32099 symptom-gene after retrieval of data (diseases and genes) from HPO database using the 255 symptoms as input. The FDR adjusted connectivity probability scores retained 239 diseases and same number of genes (3337) having 1217 and 30187 connections with 55 and 89 symptoms (total 123 symptoms) respectively. Further refining of disease-gene interactions through MalaCards scores, 134 diseases connected with 269 genes through 369 connections whereby same number of diseases (134) remained connected with 123 symptoms as mentioned above. The sequential implementation of weightages of symptoms over diseases followed by diseases over genes and their FDR adjusted connectivity probability scores retained 48 diseases and 188 genes having 265 disease-gene connections whereby 100 symptoms remained connected with selected diseases and genes. The assessment of intra-links of nodes (viz. symptom-symptom, disease-disease, gene-gene) using cosine and STRING-combined scores finally retained 329 nodes including 92 symptoms, 48 predictive diseases and predictive 189 genes with total 3114 edges to form TN (network density 0.029, average clustering co-efficient 0.108). The stepwise results obtained during TN construction are presented in the pictorial diagram of Fig. 2.

### 2. Finding ‘target nodes’ of symptoms, diseases, genes under TN

The topological assessment on TN evaluated 73 symptoms, 47 predictive diseases and 27 genes (Fig. 3a) under three different properties namely *‘both HB and driver’, ‘pure driver, ‘pure HB’* nodes. Further, the 73 important ‘target symptoms’ *(‘both HB and driver’:’pure drive’:‘pure HB*’=16:44:13) and 47 important predictive neurological ‘target Diseases’ *(‘both HB and driver’:‘pure drive’:‘pure HB’=8:0:39)* were classified into respective six and eight different categories (Fig. 3a). The 27 ‘target genes’ showed properties of *‘both HB and driver’ (CTNNB1), ‘pure HB’* (16 genes mentioned in Fig. 3d and 3E) and *‘pure driver’* nodes (10 genes mentioned in Fig. 3d and Fig. 3e).

**Fig. 3.**
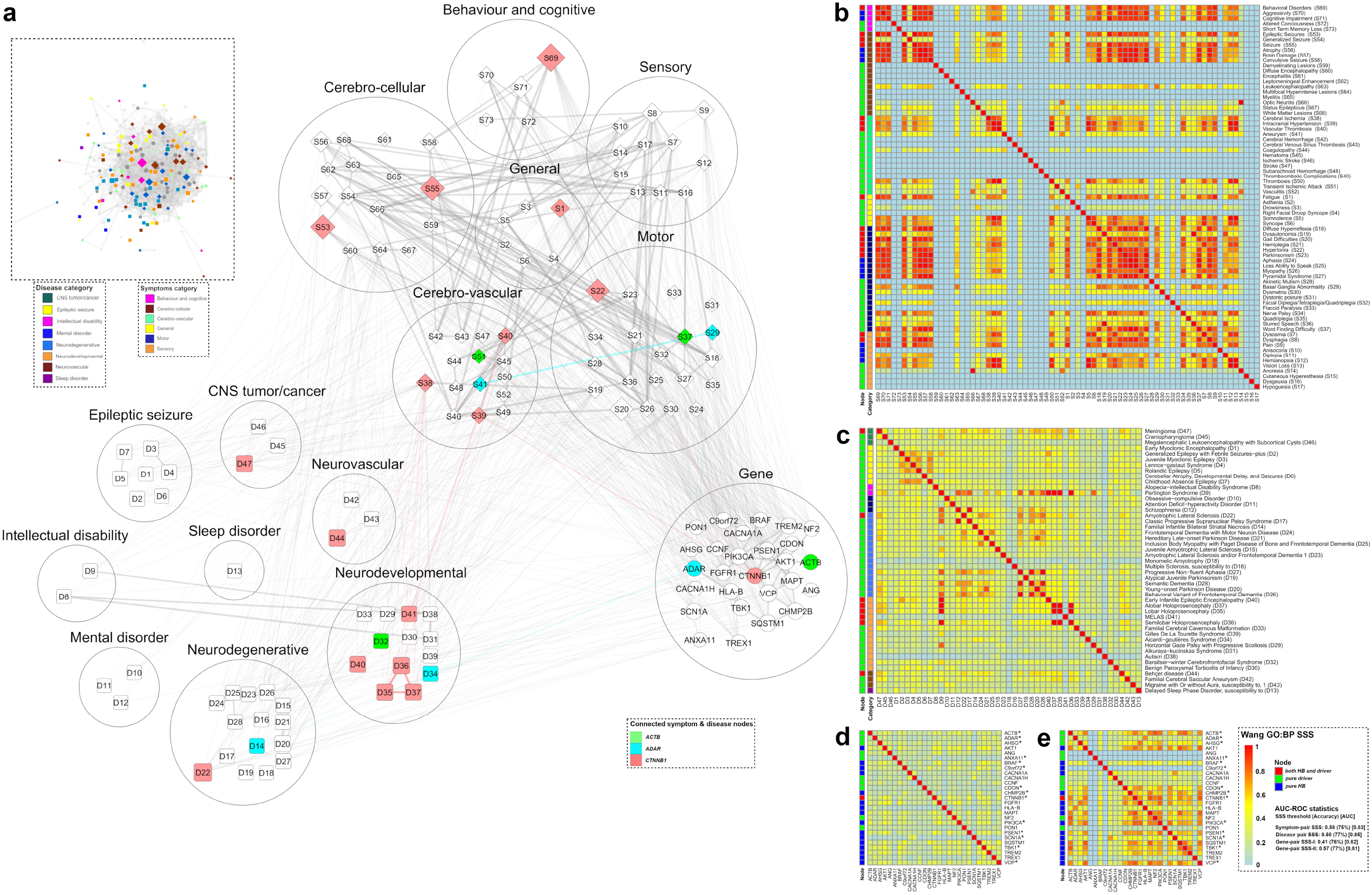
The model of ‘tripartite network’ (TN) and its *‘target nodes’* with their pairwise semantic similarity scores (SSS) for COVID-19: **(a)** The TN (inset) is developed in Cytoscape software using total 329 nodes including selected 92 symptoms of COVID-19, 48 diseases and 189 genes with their 3114 edges as described in Fig. 2. The coloured nodes (inset) represents 147 *‘targetnodes’* (HB and driver, pure driver, pure HB) including 73 symptoms, 47 diseases and 27 genes. The colour codes of nodes in TN (inset) indicate six categories of symptoms and eight categories of diseases. The pictorial presentation of ‘target nodes’ including categories (circles) of symptoms and diseases clearly indicates the shape of their connections within TN. The colour codes (light-green, cyan and light-red) in pictorial image indicate three gene nodes (ACTB, ADAR and CTNNB1) exhibiting triangular (open) inter-links with nodes in symptoms and diseases with their respective edges. Other nodes (inset and pictorial images) represented as white-colored with gray border and their connections (gray colour edges) do not show triangular inter-links. Nodes (inset and pictorial images) are represented with different shapes (diamonds for symptoms, rectangle for diseases and circle for genes) and sizes (connectivity values adjusted by the ‘continuous mapping of node size’ in ranges between 25 and 60 pts. for the lowest and highest node size respectively) having grey color borders (illustrated with 2 pts.). The widths of the edges (inset and pictorial images) are represented by respective metrics which are adjusted by 0.5 to 2 pts. of ‘continuous mapping of edge width’ in the edge network style of the Cytoscape. **(b-e)** The heat maps represent the pairwise GO-BP SSS of *‘target nodes’* of TN including 73 symptoms **(b)**, 47 diseases **(c)** and 27 genes **(d-e)**. The SSS are adjusted in continuous color gradient of light-blue:yellow:red to their respective values over the ranges of 0:0.5:1 using *pheatmap* library function in R environment. The pair-wise SSS measurements (vide ‘Methodology’ section) are calculated as *‘direct association’* (SSS-I) for genes **(d)** and *‘indirect association’* (SSS-II) for symptoms **(b)**, diseases **(c)** and genes **(e)**. The codes of symptoms **(b)** and diseases **(c)** are represented in columns of heatmaps to minimize the spaces. The vertical bars (left side of each heat map) with different colors demonstrate nodes having topological properties of centrality measurements and categories (inset in **a**) of symptoms and diseases of network. The summary of classifier ROC-AUC statistics (threshold scores, accuracy scores in %, AUC scores) of SSSs **(b-e)** are presented adjacent to the colour bar.

### 3. Reconstruction of ‘combined gene network’ (CGN) using selected co-expressed genes (CG) and genes of TN for COVID-19

#### 3.1. Finding co-expressed genes (CG) and formation of CGN

The 27 *‘target genes’* (nodes with properties of *‘both HB and driver’, ‘pure driver’, ‘pure HB’)* evaluated from TN used as query genes against a set of 38306 genes from RNA-Seq data of brain frontal cortex of COVID-19 patients in *geneRecommender* analysis and identified that 10913 genes (28.48% of total genes) ranked (functionally correlated to query genes) as co-expressed genes (CG) with the set of 14 query genes (marked ‘*’ in Fig. 3d and Fig. 3e). The selected CGs (10913 genes including 14 query genes) were further refined in Mutual Information (MI)-based network analysis using *ARACNe* algorithm in *minet* (Mutual Information NETwork inference) to find pair-wise gene connections from where a set of connected genes corresponding to each of 14 query genes had been manually curated. The 14 query genes provided an exclusive set of 225 co-expressed genes.

The total 414 genes including the exclusive 225 co-expressed genes of *minet* output and 189 genes including 27 *‘target genes’* found in TN were used as input for STRING database and provided a PPI (Protein-Protein Interaction) network (network density 0.010, average clustering co-efficient 0.169) which was designated as *‘combined gene network’* (CGN) of 281 gene nodes, using SPPICS confidence threshold values >0.60. (Fig. 4b)

**Fig. 4.**
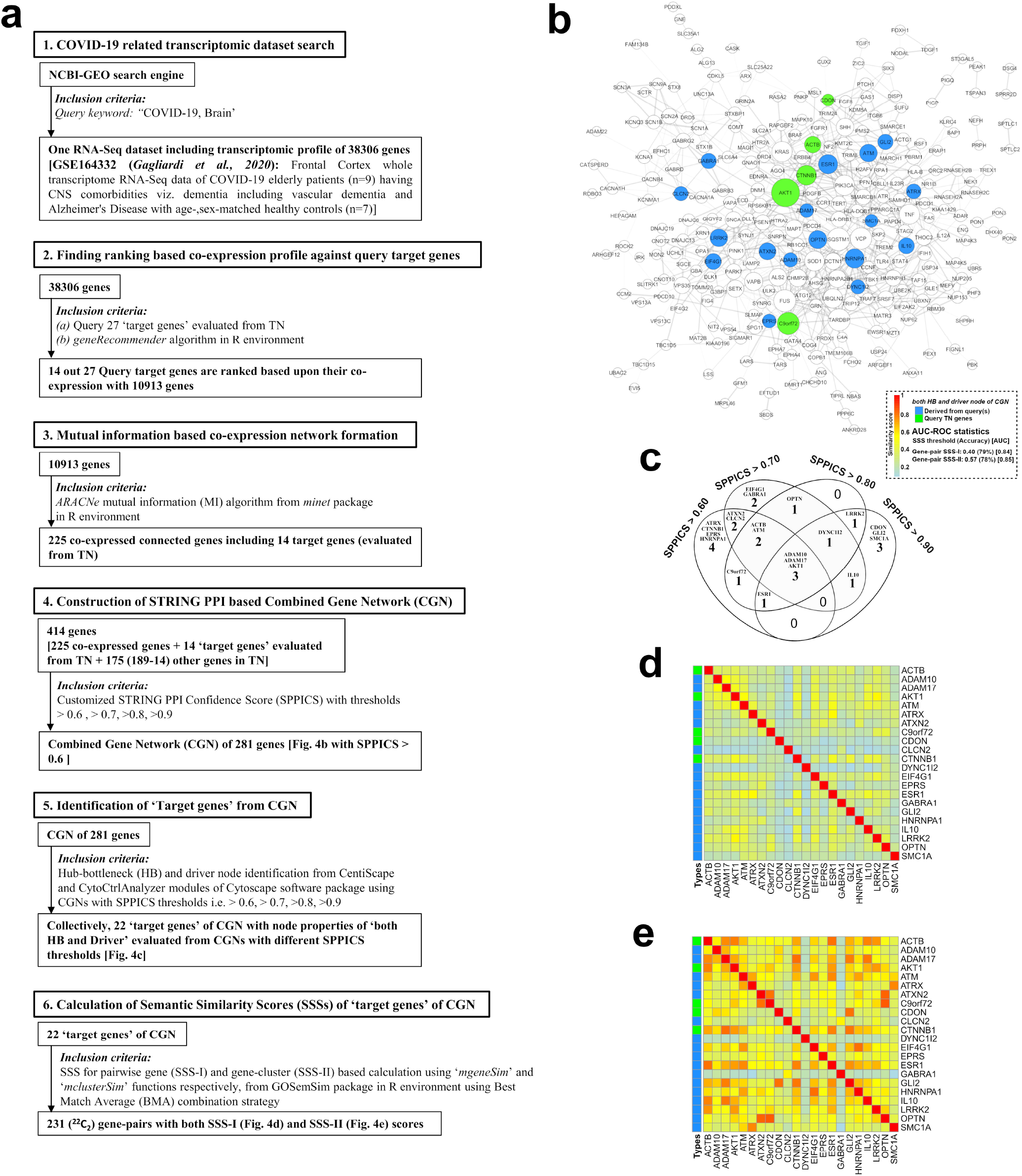
The model of ‘combined gene network’ (CGN) and its *‘target nodes’* with their pairwise semantic similarity scores (SSS) for COVID-19: **(a)** Study design for the construction of the CGN is represented stepwise (point 1-6). **(b)** The PPI interactome model of CGN is a continuous network consisting of total 281 nodes of gene products/proteins and 793 edges corresponding to the functional connectivity between nodes. The CGN is constructed using SPPICS >0.60 as widths of the edges which are adjusted by 0.5 to 5 pts. of ‘continuous mapping of edge width’ in the edge network style of the Cytoscape. The colour nodes (inset) represent 22 *‘target genes’* having both ‘HB and driver’ properties including five ‘query genes’ derived from TN (green nodes) and 17 ‘co-expressed genes’ (dodger blue color) derived from mRNA-Seq data of COVID-19 patients. The other nodes of the CGN are kept white-colored with grey colored border (illustrated with 2 pts.). The sizes of the nodes indicate their connectivity (higher the value, higher will be the size) adjusted by the ‘continuous mapping of node size’ in ranges between 25 and 60 pts. for the lowest and highest node size respectively. (**c**) The Venn diagrams indicate the changes in number of ‘target genes’ (both ‘HB and driver’) with their names derived from CGNs developed using multiple SPPICS thresholds including >0.60, >0.70, >0.80 and >0.90 for inclusion of all possible ‘target genes’ for better interpretations. (**d and e)** The heat maps represent the pairwise GO-BP SSS of 22 *‘targetgenes’* of CGN. The colour codes (light-blue:yellow:red) of the values of SSS in heat maps range 0:0.5:1 developed in *pheatmap* library function in R environment. The values of SSS-I (**d**) and SSS-II (**e**) are direct and indirect associations (vide ‘Methodology’ section) respectively. All ‘target genes’ in CGN belong to ‘HB and driver’ topological properties of centrality measurements. The colours of the vertical bars (left side of each heat map) indicate types (vide inset) of *‘target genes’.* The summary of classifier ROC-AUC statistics (threshold scores, accuracy scores in %, AUC scores) of SSSs are presented in adjacent to the colour bar.

#### 3.2. Evaluation of ‘target genes under CGN’ (TG-CGN)

The study was extended to construct STRING-PPI networks using same set of genes as input with different SPPICS confidence threshold score viz. >0.70, >0.80 and >0.90 for obtaining sets of ‘target genes’ from different PPI networks. The topological centrality and controllability studies on four separate CGNs provided a set of 13, 12, 10 and 10 *‘both HB and driver’* nodes (Fig. 4c). Collectively, the 22 gene nodes were obtained as *‘both HB and driver’* from the four different PPI-networks using different confidence score (SPPICS) threshold values and designated as *‘target genes under CGN’* (TG-CGN) for COVID-19 (Fig. 4c). Interestingly, five genes such as *ACTB, AKT1, C9orf72, CDON* and *CTNNB1,* part of 14 query genes evaluated from TN, appeared to be included in the set of 22 TG-CGN (green colored nodes in Fig. 4b).

## 4. Evaluation of pair-wise values of SSS for ‘target nodes’

### 4.1. Finding pair-wise SSS-II values of ‘target nodes’ for symptoms and diseases evaluated from TN

The 73 symptoms and 47 diseases as ‘target nodes’ evaluated from TN, provided pairwise SSS-II scores for 2628 (^73^C_2_) and 1081 (^47^C_2_) combinations (Fig. 3b and Fig. 3c). The 45.53% symptom-pairs and 12.01% diseases-pair had been found significant based on the ‘optimal threshold for ROC’ as cut-off values.

The categorical investigation of filtered symptom-pairs resulted 22.42% intra-group and 77.57% inter-group connections including the categories (%inter:%intra) under (i) general (9.40:1.67), (ii) sensory (22.41:9.17), (iii) motor (32.53:69.17), (iv) cerebro-vascular (13.01:5.83), (v) cerebro-cellular (14.94:11.67) and (vi) behaviour and cognitive (7.71:2.50) (Fig. 3b). In case of filtered disease-disease pair SSS, 36.15% intra- and 63.85% inter-group connectivity were observed, including (i) epileptic seizure (1.20:21.28), (ii) intellectual disability (7.23:0.00), (iii) mental disorder (4.82:2.13), (iv) sleep disorder (0.00:0.00), (v) neurodegenerative (12.05:61.70), (vi) neurodevelopmental (51.81:12.77), (vii) neurovascular (7.23:0.00) and (viii) CNS tumour/cancer (15.66:2.13) (Fig. 3c).

### 4.2. Finding pair-wise values of SSS-I and SSS-II of ‘target genes’

The 27 TG-TN showed 351 (^27^C_2_) gene-pairs with SSS-I values whereas provided 276 (^24^C_2_) gene-pairs with SSS-II values, because, three gene nodes *(ANG, ANXA* and *C9orf72*) did not have PPI connections in TN and SSS-II was calculated based on connected cluster of genes. The 22 TG-CGN genes exhibited 231 (^22^C_2_) gene-pairs with both SSS-I and SSS-II values providing that the 22 genes had connections with other genes in CGN. Irrespective of the networks (TN and CGN) all gene-pairs showed same SSS-I values and SPPICS values also appeared same for respective gene-pairs (Fig. 3d and Fig. 4d). Further refining based on the ‘optimal threshold for ROC’ as cut-off values, the SSS-I scores resulted statistically significant 31 and 33 gene-pairs respectively for TG-TN and TG-CGN. Similarly, the SSS-II scores (Fig. 3e and Fig. 4e) resulted statistically significant 81 and 143 gene-pairs respectively for TG-TN and TG-CGN.

Among selected gene-pairs in TN based on significant SSS-I (31 gene-pairs, Fig. 3d) and SSS-II (81 gene-pairs, Fig. 3d) values, 20 gene-pairs found to be common in the set (Fig. 3d and 3e) and nine of common gene-pairs consisted physical PPI interactions with SPPICS values in the TN (Fig. 3a). The nine common gene-pairs having SSS-I, SSS-II and SPPICS values are *FGFR1-PIK3CA* (0.535,0.809,0.97)*, SQSTM1-VCP* (0.513,0.785,0.704)*, AKT1-CTNNB1* (0.487,0.783,0.997)*, CHMP2B-SQSTM1* (0.484,0.729,0.81)*, AKT1-PIK3CA* (0.482,0.79,0.998)*, CTNNB1-PSEN1* (0.469,0.68,0.994)*, MAPT-PSEN1* (0.454,0.64,0.885)*, AKT1-MAPT* (0.451,0.729,0.804) and *AKT1-VCP* (0.419,0.798,0.776). Similarly, out of the 33 (Fig. 4d) and 143 (Fig. 4e) gene-pairs of CGN having SSS-I and SSS-II values respectively, 33 gene-pairs found to be common in the set and among them 13 gene-pairs showed physical PPI interactions with SPPICS values in the CGN (Fig. 4b). The thirteen common gene-pairs having SSS-I, SSS-II and SPPICS values are *ADAM10-ADAM17* (0.558,0.771,0.863), *AKT1-ATM* (0.535,0.714,0.804), *ADAM17-AKT1* (0.496,0.784,0.735), *AKT1-CTNNB1* (0.487,0.805,0.997)*, ESR1-CTNNB1* (0.483,0.833,0.842), *AKT1-EIF4G1* (0.475,0.666,0.645), *AKT1-ESR1* (0.467,0.803,0.991), *AKT1-IL10* (0.461,0.773,0.828), *GLI2-CTNNB1* (0.45,0.851,0.613), *ATM-ATRX* (0.443,0.788,0.713), *LRRK2-CTNNB1* (0.443,0.702,0.834), *C9orf72-OPTN* (0.428,0.885,0.815), *C9orf72-ATXN2* (0.402,0.856,0.754). Interestingly, the PPI link viz. *AKT1-CTNNB1* was evident both in TN and CGN (Fig. 3a and 4b). Therefore, total 21 gene-pairs including eight from TN, 12 from CGN and one common for both TN and CGN found to have SSS-I, SSS-II and SPPICS values and, were designated as *‘candidate gene pairs’* for COVID-19 (Fig. 5f).

**Fig. 5.**
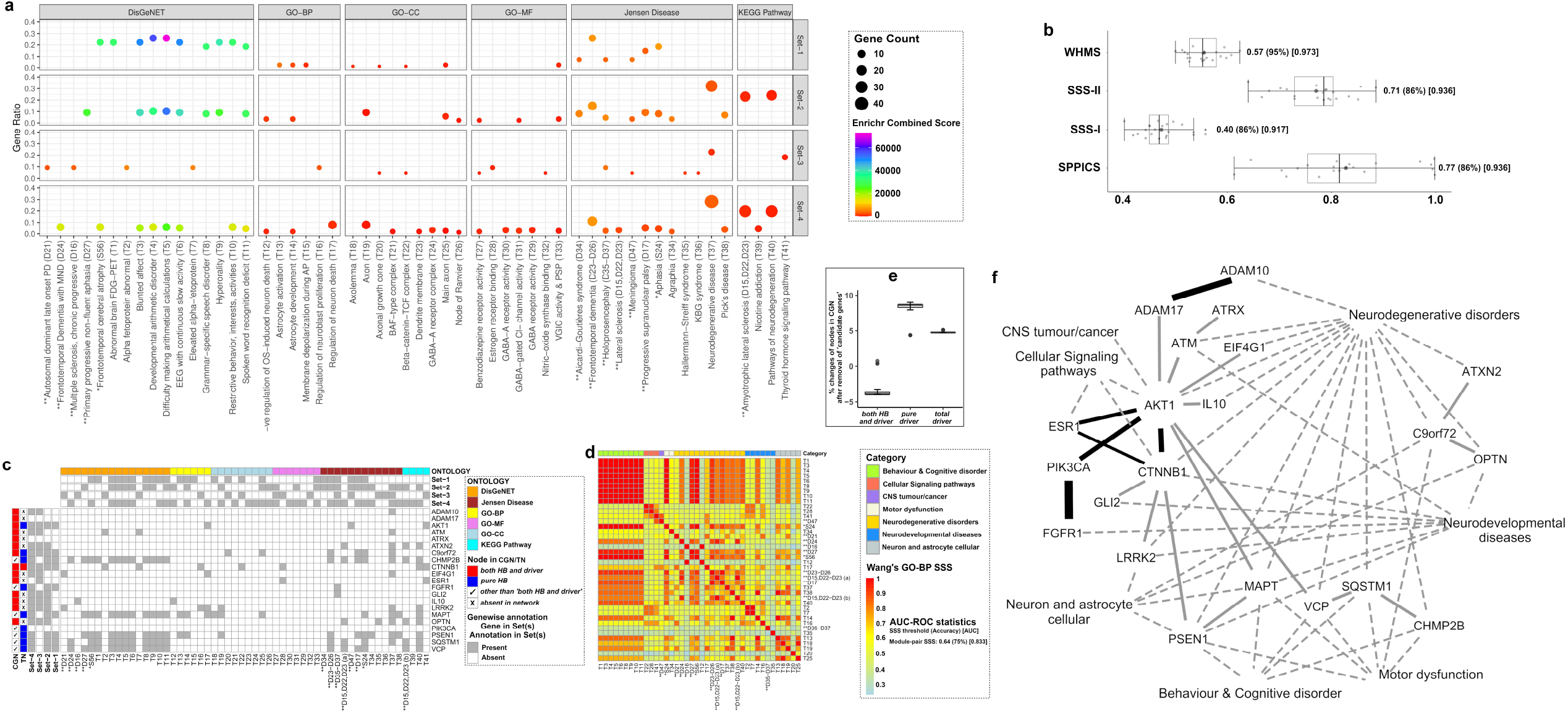
The ‘Candidate genes’ and their functional modules for neurological manifestations of COVID-19: **(a)** The bubble plot exhibiting the results of Enrichment analysis. The statistically significant (FDR adjusted p-value <0.05) 54 “terms” (X-axis) specific for nervous system found under Enrichr functional annotations (GO terms, KEGG pathways, DisGeNET and Jensen Diseases) are identified by ‘target genes’ derived from TN (set-1) and CGN (set-3) and their connecting genes (Set-2 for TN and Set-4 for CGN) in corresponding networks. Each term is characterized by ‘gene count’ (size of the bubble adjusted with 0.5 to 4 pts.) and corresponding ‘gene ratio’ (Y-axis). All terms showed ‘combined score’ (log(p-value) × z-value) with most stringent values (>147) which are adjusted in a color (VIBGYOR) gradient. (**b**) The box plot of the data (X-axis) of multiple interaction scores (SPPICS, SSS-I, SSS-II and WHMS in Y-axis) of 21 selected pairs of ‘candidate genes’, constructed by *geom_boxplot()* function of *ggplot2* in R package. The pair of ‘candidate genes’ are selected from pairs of ‘target genes’ of TN and CGN which have all three values of SSS-I, SSS-II and SPPICS; they were considered for evaluation of integrated WHMS. (**c**) The checkerboard to demonstrate the abundance (presence/absence as gray/white colour) of 21 ‘candidate genes’ associated with enriched 54 functional annotation ontology terms manually curated from results of the bubble plot (A). The vertical bars (left side) represent the status of genes as ‘both HB and driver’ (red clour), ‘pure HB’ (blue colour), other than ‘both HB and driver’ (tick mark as ‘pure driver’ and ‘non-HB non-driver’) in TN/CGN. The horizontal bars (at top) represent the enrichment terms with ontology categories (six different colours) and their presence/absence (gray/white colour) in sets for enricher analysis (collected from A). **(d)** The heat map to represent the pairwise GO-BP SSS-II of 40 enriched terms overrepresented with at least one candidate genes. The SSS-II range from 0.23 to 1 (colour light-blue:yellow:red). The horizontal bar (at top) represents categories of enriched terms (seven different colours). **(e)** The boxplot to represent the % changes of ‘target nodes’ in CGN after removing one ‘candidate gene’ by leave-one-out method to cross-validate disease-causing ‘*indispensable*’ driver node property. **(f)** The pictorial presentation (developed in Cytoscape software) of interactome model of 21 ‘*candidate genes*’ and their associated functional seven categories of Enrichr terms related to neurological manifestations of COVID-19. The solid lines (width adjusted by 3 to 10 pts. based on WHMS) represent gene-pair interactions with the ‘prevalent’ links (dark solid lines) of *‘candidate genes’* which are selected based on WHMS more than cut-off values (‘optimal threshold score of ROC’ given in **e**). The dotted lines (width adjusted by 2 pts.) represent links between gene and functional categories of Enrichr terms. The boxplots (**b** and **e**) represent categorically, the quartile values (first quartile:Q1 as left edge of box; third quartile:Q3 as right edge of box), inter quartile ranges (IQR), median values (black vertical line inside box), mean values (grey coloured filled circles), maxima (Q3+1.5*IQR)/minima (Q1-1.5*IQR) (two vertical lines) and their out layers of data. The classifier statistics ROC-AUC summaries (optimal threshold score, accuracy score in % and AUC score) are presented in respective cases (right side of boxplots in **b**; adjacent to the colour bar in **d**). The codes (**5c** and **5d**) ‘D15,D22,D23(a)’ and ‘D15,D22,D23(b)’ represent Enrichr annotation terms as ‘Lateral sclerosis’ (ONTOLOGY: Jensen Disease) and ‘Amyotrophic lateral sclerosis’ (ONTOLOGY: KEGG pathway) respectively.

### 5. Selection and categorization of ‘candidate gene’ *pairs* by the formulation of WHMS

The selected 21 *‘candidate genes’* with pair-wise 21 interactions *(‘candidate gene pair’)* showed excellent (AUC>0.90) classifications (sets) based on interaction values of SSS-I, SSS-II and SPPICS. The SSS-I, SSS-II and SPPICS values showed same accuracy score as 0.86 and that was used as weightage for respective case to calculate WHMS of 21 pair-wise interactions of the *‘candidate genes’.* The integrated values of WHMS exhibited accurate (95%) and excellent classifications with highest AUC value (0.97) of those 21 gene-pairs and therefore the values of WMHS considered as better choice of further analysis (Fig. 5b and Table 1).

**Table 1.**
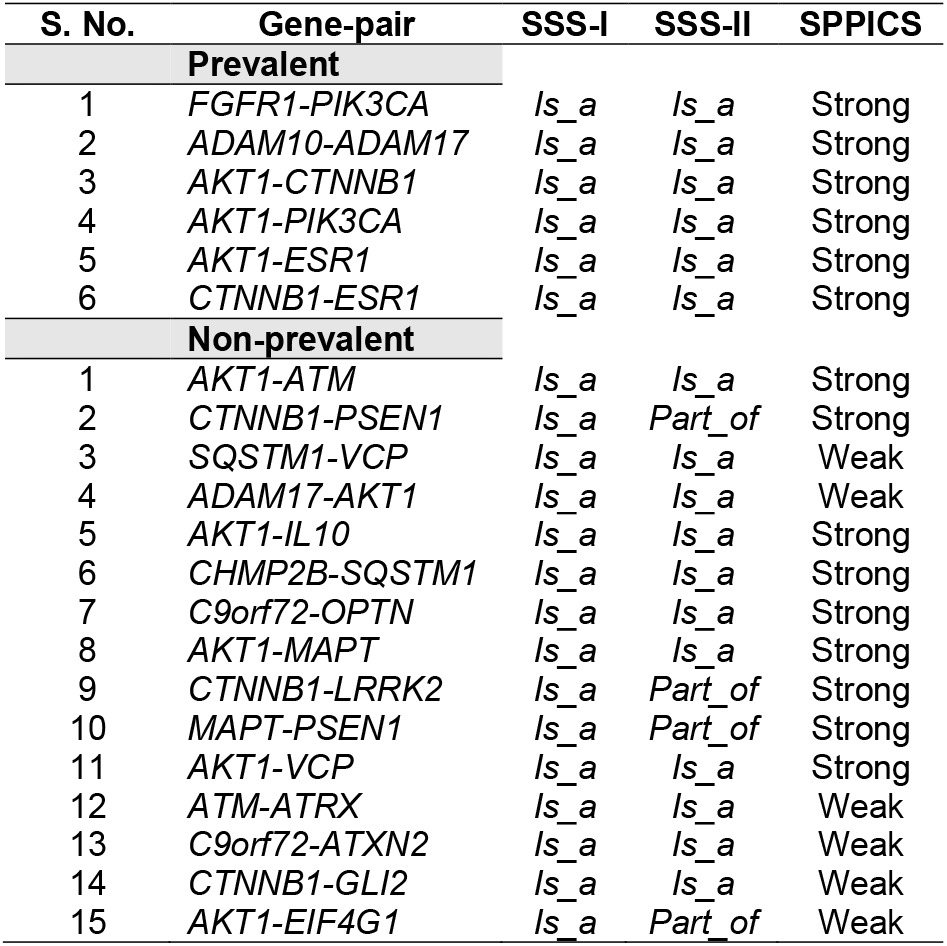
Summary of pair-wise ‘candidate genes’ with their properties evaluated as ‘prevalent’ and ‘non-prevalent’ characters and their functional links having statuses with SSS-I, SSS-II and SPPICS values. The prevalent pair-wise ‘candidate genes’ are selected based on values of gene-pairs more than cut-off value of WHMS (vide Fig. 5B & 5F). The terms *‘Is_a’* (subtype) *‘Part_of”* (component) represent the functional associations of GO based terms hierarchically (ancestors-descendants relationship) organized in directed acyclic graph (DAG). The terms ‘*Is_a*’ and ‘*Part_of*’’ properties are considered to the gene-pairs with scores greater than and less than cut-off values of SSS-I and SSS-II respectively. The terms ‘Strong’ and ‘Weak’ represent the link strength having values of SPPICS to gene-pairs with scores greater than and less than cut-off value respectively. The cut-off values (optimal threshold score of ROC) are given in Fig. 5B.

The six pair-wise interactions of *‘candidate genes’* viz. *ADAM10-ADAM17* (WHMS: 0.61), *AKT1-CTNNB1* (WHMS:0.60), *AKT1-ESR1* (WHMS:0.59), *AKT1-PIK3CA* (WHMS:0.59), *CTNNB1-ESR1* (WHMS:0.58) and *FGFR1-PIK3CA* (WHMS:0.62) showed WHMS values greater than the cut-off value i.e., ‘optimal threshold value of ROC’ (0.57) for WHMS (Fig. 5b) and considered them as prevalent pairs-wise interactions of the *‘candidate genes’* for discussion of their functional aspects in COVID-19 (dark solid edges in Fig. 5f). Notably, six pair-wise prevalent *‘candidate genes’* had values of SSS-I, SSS-II and SPPICS greater than their respective cut-off values i.e., ‘optimal threshold values of ROC’ (SSS-I:0.40, SSS-II:0.71, SPPICS:0.77) (Fig. 5b and Table 1).

The 15 pair-wise interactions of *‘candidate genes’* showed WHMS values below the cutoff value (light solid edges in Fig. 5f). These 15 interactions appeared to have values of SSS-I, SSS-II and SPPICS greater than respective cut-off values except few cases with low values of SSS-II (four gene pairs viz. *CTNNB1-PSEN1*, *CTNNB1-LRRK2, MAPT-PSEN1, AKT1-EIF4G1)* and SPPICS (six gene pairs viz. *SQSTM1-VCP, ADAM17-AKT1, ATM-ATRX, C9orf72-ATXN2, CTNNB1-GLI2, AKT1-EIF4G1).* In fact, all 21 pair-wise ‘*candidate genes*’ showed SSS-I values greater than its cut-off values (Table 1).

### 6. Enrichr analysis of ‘target genes’ and Integration of annotation terms for COVID-19

Enrichr annotation of four sets of genes having *‘target genes’* (set-1 and set-3) and their connected genes (Set-2 and set-4, respectively) resulted statistically significant total 159 terms (data not shown) including several common terms among four sets. The statistically significant (FDR adjusted p-value <0.05) Enrichr annotation terms (total 159 terms) were further cross-validated manually with selected (above cut-off values of SSS-II depicted in Fig. 3b and Fig. 3c) *‘target nodes’* (HB and driver) of the symptoms and diseases found in TN analysis (Fig. 3a). 13 common terms including two symptoms and 11 diseases appeared to distribute (Fig. 5a) in (i) DisGeNet annotation (four diseases and one symptom), (ii) Jensen diseases annotation (six diseases and one symptom) (iii) KEGG pathway annotation (one disease). Furthermore, the rest of the total terms (146, data not shown) showed selected 41 terms associated with functional annotation of nervous system including 11 in DisGeNET (T1-T11), six in GO-BP (T12-T17), nine in GO-CC (T18-T26), seven in GO-MF (T27-T33), five in Jensen Disease (T34-T38) and three in KEGG pathway (T39-T41) were selected for functional integration with genes among four sets (Fig. 5a). Therefore, 54 (41 + 13) selected terms were implied to find their links with candidate genes in subsequent analysis.

### 7. Finding functional annotations of ‘candidate genes’ and their categorization for COVID-19

Total 40 functional annotation terms had been found for 21 *‘candidate genes’* represented in the checkerboard diagram (Fig. 5c). The selected 40 functional annotation terms were categorized in to six different functional modules viz. ‘Behaviour & Cognitive disorder’, ‘Cellular Signaling pathways’, ‘CNS tumour/cancer’, ‘Motor dysfunction’, ‘Neurodegenerative disorders’, ‘Neurodevelopmental diseases’, ‘Neuron and astrocyte cellular’, comparing with functional modules categorized for symptoms and diseases in TN (Fig. 3b and Fig. 3c).

The semantic similarity assessment provided 780 pair-wise (^40^C_2_) SSS-II values of 40 functional annotation terms using their connected ‘*candidate genes*’. The SSS-II values for pair-wise 40 functional terms showed an accurate (75%) and good classification (AUC:0.83) with 330 SSS-II values above the ‘optimum threshold of ROC’ i.e., cut-off value (0.64) having 28.78% and 71.21% of terms under intra- and inter-category (module) similarity, respectively (Fig. 5d). The terms (54.89%) under ‘Behaviour & Cognitive disorder’ showed highest inter-category similarities with terms of the categories under ‘Cellular Signaling pathways’ (2.32% terms), ‘Motor dysfunction’ (6.97% terms), ‘Neurodegenerative disorders’ (55.81% terms), ‘Neurodevelopmental diseases’ (6.97% terms) and ‘Neuron and astrocyte cellular’ (27.9% terms). The terms (51.58%) under neurodegenerative disorders showed highest intra-category similarities among terms (Fig. 5d).

Interestingly, certain pair-wise terms appeared common among statistically significant terms under (a) enriched for ‘*candidate genes*’ (Fig. 5d) and (b) ‘*target nodes*’ for symptoms/diseases under TN (Fig. 3a) and, found to have greater pair-wise SSS-II values with ‘*candidate genes*’ viz. atrophy-aphasia (3.7% more SSS-II value) as symptom-pair (Fig. 3b and Fig. 5d) and other five disease-pairs including (a) ‘progressive non-fluent aphasia’ with ‘classic progressive supranuclear palsy syndrome’ (15% more SSS-II value), ‘amyotrophic lateral sclerosis’ (32.5% more SSS-II value), ‘frontotemporal dementia’ (17.7% more SSS-II value) and (b) ‘frontotemporal dementia’ with ‘amyotrophic lateral sclerosis’ (8.6% more SSS-II value) and ‘progressive supranuclear palsy syndrome’ (17.2% more SSS-II value) (Fig. 3c and Fig. 5d).

### 8. Finding essentially of *‘candidate genes’*

Cross-validation through leave-one-out method of 21 candidate-gene from the 281 gene node containing CGN network resulted 3.15% average reduction in *‘both HB and driver* node and average increase in 7.96% and 4.81% *‘pure driver’* and *‘driver’* nodes, respectively (Fig. 5e). The results signified that the 21 ‘*candidate genes*’ appeared to be *‘indispensable’* driver nodes and supposed to be COVID-19 predictive neurological disease-related regulatory genes.

### 9. Assimilation of *‘candidate genes’* with functional annotations to develop a brain related functional modules for COVID-19

The 21 *‘candidate genes’,* their 21 interactions having WHMS and links with corresponding functional annotations had been integrated in pictorial diagram (Fig. 5f) for better interpretations. The seven *prevalent ‘candidate genes’ (ADAM10, ADAM17, AKT1, CTNNB1, ESR1, FGFR1, PIK3CA)* showed direct association with enriched terms under the categories of ‘Cellular Signaling pathways’, ‘CNS tumour/cancer’, ‘Neurodegenerative disorders’ and ‘Neurodevelopmental diseases’. Their indirect association with enriched terms under the categories of ‘Neuron and astrocyte cellular’ events, ‘Behaviour & Cognitive disorder’ and ‘Motor dysfunction’ appeared through interactions with other (non-prevalent) *‘candidate genes’* (Fig. 5f).

## Discussion

The present study is a novel approach of combined meta-analysis of two networks viz. TN and CGN to find the *‘disease-related regulatory genes’* associated with functional (transcriptional and translational) cellular entities necessary for understanding molecular basis of brain pathophysiological phenotypes of COVID-19. To achieve the goal, we proceeded with most robust approaches through multiple screening steps including (a) finding the two sets of predictive ‘*target genes*’, evaluated from TN and CGN with their PPIs having STRING *‘combined scores’* (SPPICS) as priori analysis, (b) evaluation of functional associations by ‘semantic similarity scores’ (SSS) of two sets of *‘target genes’,* (c) screening *‘target genes’* by cumulating PPIs having both STRING-CS and SSS by selection with given threshold value for respective PPI scores to find ‘candidate genes’, (d) formulating integrated scores (WHMS) combining SPPICS and SSS for giving weightage to PPIs of *‘candidate genes’* for further categorization, (e) assimilation of ‘*annotation terms*’ (symptoms/diseases) with genes among ‘*candidate nodes*’ through posteriori enrichment analysis to get functional module. Notably, ‘*target nodes*’ for symptoms and diseases evaluated from TN were manually curated and integrated with suitable Enrichr annotations for better interpretation. The classification statistics (ROC-AUC) and cut-off values (optimal threshold for ROC) identified the PPIs with their association scores (SPPICS, SSS, WHMS) for respective steps most accurately (AUC>0.8) with minimum false positive interpretation. Furthermore, all 21 *‘candidate genes’* appeared to be almost equally *‘indispensable’* after screening of controllability property of ‘candidate genes’ on CGN. Finally, the ‘*candidate genes*’ were categorized based on pair-wise analysis of values of WHMS, SSS and SPPICS to find prevalent vs. non-prevalent *‘candidate genes’* with their pattern *(‘is_a’* vs *‘part_of’)* of relationship with neurological manifestation in COVID-19. The pathophysiological relevance of prevalent *‘candidate genes’* with COVID-19 have discussed thoroughly.

In the present study, two networks (TN (Fig. 3a) and CGN (Fig. 4b)) were analyzed to find the *‘target nodes’* which satisfied the three properties viz. *‘hub’, ‘bottlenecks’* and *‘driver’* together for COVID-19. Separate studies indicates that host proteins targeted by viral proteins show the node properties of hubs and high betweenness centrality^82^ and, *‘indispensable’* driver controllability^69,71^ in a host protein network. The *‘target genes’* (TG) evaluated from TN (TG-TN) showed node properties of hub-bottleneck (HB or *‘date-hubs’* i.e., together hub and bottlenecks), driver and both (HB and driver) (Fig. 3d and Fig. 3e). All *‘target genes’* (TG) evaluated from CGN (TG-CGN) showed node properties as both HB and driver (Fig. 4b and Fig. 4c). In fact, the number of driver nodes compared to the driver nodes themselves appears as crucial for maintenance the controllability of a network^69,71^. In the present study, the finally selected 21 *‘candidate genes’* appeared as *‘indispensable’* as the number of driver nodes increased (4.81 % for total driver, 7.96% for driver but non-hub-bottlenecks i.e., pure driver) in the CGN after removal of one of the ‘*candidate genes*’ (Fig. 5e).

The SPPICS were applied to construct possible PPI connections of new genes in TN (Fig. 3a) and CGN (Fig. 4b) networks related to brain in COVID-19. The SPPICS provides quantitative measurement of physical and functional PPI evidences derived from available online resources. It is devoid of experimental evidences of functional entities related to regulatory mechanism (transcriptional and translational regulation) in physiological context of cells, as part of its calculation^56^. The Gene Ontology resources provides a model of hierarchically (ancestors-descendants relationship) organized directed acyclic graph (DAG) having GO-terms as nodes and functional association as directed edges within each hierarchy by ‘*is_a*’ (subtype) and ‘*part_of*’ (component) relationship associated to gene/protein functionality (molecular function, cellular component and biological process) description^73,74^. GO based biological process (GO-BP) provides cohesive evidences about protein interactions related to both physical and functional networks of molecular events involved in cellular physiology^83–85^. The candidate genes for a disease show common biological pathway(s)^48^. Therefore, in the present study, the functional association among *‘target nodes’* were analyzed by semantic comparison of GO-BP annotations quantitatively through computing similarities between gene-pairs (SSS-I measurement) (Fig. 3d and Fig. 4d) and also clustering gene/symptoms/disease/module-pairs (SSS-II measurement) into known pathways (Fig. 3b, Fig. 3c, Fig. 3e, Fig. 4e and Fig. 5d).

The conventional SSS-I provided pair-wise ‘*direct association*’ based on comparative assessment of associated GO-BP terms of two target genes (Fig. 3d and Fig. 4d). It is reported that the genes and their functionally connected co-expressive genes show tissue specific expressions and regulations (transcriptional and translational) and, exhibit pleotropic effects i.e., share common symptoms and diseases^86–89^. Based on this concept, the estimation of SSS-II values was newly introduced in the present study (Fig. 3e and Fig. 4e). The SSS-II values provided pair-wise *‘indirect association’* based on the summated contribution of comparative assessment of associated GO-BP terms of geneclusters (connected genes) against targeted gene-pairs. Our data indicated that classification of both SSS-I and SSS-II values were statistically robust (AUC: 0.91 and 0.93) with the different range of values and had respective accurate (0.40 and 0.71) threshold values for ROC to interpret the results in most stringent way (Fig. 5b). Notably, the SSS-I and SSS-II values for gene-pairs represented very interesting observations to interpret the results. The gene-pairs found as common PPI both in TN and CGN networks showed same values of SSS-I whereas SSS-II values varied for networks. For example, *CTNNB1*-*AKT1* gene-pair among ‘*target genes*’, found as common PPI both in TN (Fig. 3a) and CGN (Fig. 4b), showed equal SSS-I value (0.487) (Fig. 3d and Fig. 4d). The SSS-II values of this gene-pair varied for TN (0.783) and CGN (0.805) (Fig. 3e and Fig. 4e). Additionally, certain gene-pairs having considerable (above threshold value) SSS-I values appeared to have low (below threshold value) or zero (‘null functional similarity’) SSS-II values including *AKT1-FGFR1* (SSS-I: 0.485 – above threshold; (Fig. 3D), SSS-II: 0.69 – below threshold; (Fig. 3e)), *C9orf72-SQSTM1* (SSS-I: 0.462 – above threshold; (Fig. 3D), SSS-II: 0; (Fig. 3e)). Therefore, the gene-pairs having significant values of both SSS-I and SSS-II had been considered for better interpretation of results in the present study.

Irrespective to the network, the SSS-I values of gene-pairs/PPIs might depict the global and existing *‘is_a’* and/or *‘part_of’* semantic similarity available in the GO-BP annotation data and therefore would remain same for representing generalized pathophysiological functions for any disease condition. Alternatively, the SSS-II values for gene-pairs varied due to different constituents in ‘gene-clusters’ which provided the *‘is_a’* and/or *‘part_of* functional relationship by sharing common GO-BP annotation terms to reflect the discrete or pleotropic effects of genes among networks (Table 1). Particularly, the zero value of SSS-II of a gene-pair indicated that *‘gene-clusters’* (connected genes) against the genepair did not well-supported by current literature-based evidences related to COVID-19 neurological symptoms. Therefore, the SSS-II values might provide a better disease-specific metric for the event of disassembly in the homeostatic genetic connectivity that being perturbed during COVID-19 insult.

The increased network quality enhances the prediction accuracy of finding ‘*candidate genes’* for a disease in PPI network-based model. The PPI model of STRING-database includes genes from prior knowledge and therefore has its own limitation^56^. The PPI model developed by semantic similarity scores has GO annotation biasness and unable to include genes not having sufficient annotation information. The integration of networks is supposed to perform better by filtering out the false positive interactions^81,90^. An approach has been reported to evaluate integrated score by combining semantic similarity scores of anatomy-based gene network and STRING-based PPI network by introducing *‘accuracy values of ROC’* as weightages to the respective scores followed by summation of weighted scores^81^. The same principal of weightage *(‘accuracy values of ROC)* was applied in the present study and the integrated scores (WHMS) were evaluated using harmonic mean of weightage scores for those gene-pairs which appeared to fulfil the criteria of having (a) three individual scores (SPPICS, SSS-I, SSS-II) and (b) at least one score with value above respective threshold level (Fig. 5b). Total 21 gene-pairs (solid edges in Fig. 5f) having integrated scores provided excellently strong fitted (AUC>0.9) and most accurate (95%) interactions (Fig. 5b) for prediction of 21 *‘candidate genes’* (Fig. 5c and Fig. 5f) involved in neurological insults (Fig. 5f) in COVID-19.

All 21 gene-pairs/PPIs of ‘*candidate genes*’ showed SSS-I values (Fig. 3d, Fig. 4d, vide Point 4.2 in results section) above the respective threshold value and therefore represented as *‘is_a’* functional relationship (Table 1) in the semantic similarity of GO-BP annotations for generalized pathophysiological functions irrespective of disease. Based on the threshold value of integrated PPI scores (WHMS), 21 pair-wise ‘*candidate genes*’ were classified as *‘prevalent’* and ‘*non-prevalent’ ‘candidate genes’* (Table 1). Six pairs of seven ‘*prevalent*’ ‘*candidate genes*’ showed strong database-dependent putative interaction scores (SPPICS) (Fig. 3a and Fig. 4b) and subsequently satisfied SSS-II values (Fig. 3e and Fig. 4e) above the threshold levels representing ‘*is_a*’ relationship (Table 1) with neuro-pathological manifestations in COVID-19. The ‘*non-prevalent’ ‘candidate genes’* found to have varied SPPICS scores (strong and weak) and different relationships (‘*is_a*’ and ‘*part-of*’) among their gene-pairs (Table 1). The ‘*prevalent*’ *‘candidate genes’ (ADAM10, ADAM17, AKT1, CTNNB1, ESR1, FGFR1, PIK3CA)* might have most prominent pathophysiological relevance in COVID-19.

The pathophysiological action of SARS-CoV-2 in brain tissue cells begins with its binding to ACE2 receptors of the cell membrane. After viral endocytosis is over, ADAM17 directs the shedding of the ectodomain of the receptors^91^, and enhances the formation of TNF-α with escalation of the cytokine storm^3^. Dysfunction of ADAMs can also exacerbate Alzheimer’s disease condition through the misfolded Aβ pathology^92^, ischemic stroke^93^ and vascular thrombosis^94^ via ACE2 and TNF-α receptors. Recently, *ADAM10* and *ADAM17* have been marked as the risk factors for cerebral infarction and hippocampal sclerosis related epilepsy^95^, respectively. In diabetic patients, an elevated activity of ADAM17 is found to enhance COVID-19 susceptibility^96^ through AKT1*-*mediated pathway.

*AKT1* encodes protein kinase B, which is a part of PI3K-NFκB signaling pathway, involved in aberrant expression of IL-10 and inflammation in severe coronavirus infection^92^. AKT1 can induce tumor formation through upregulation of RNA binding protein EIF4G1^97^, coronavirus exit from endosomes via valosin containing protein VCP^98^ and MAPT-associated tau protein formation in dementia like cognitive impairment^99^. The altered AKT1 signaling pathway is also evident in ATM-associated autism spectrum disorder that may exaggerate in COVID-19^100^.

*CTNNB1* expresses β-catenin related to Wnt signaling pathway and gets downregulated in COVID-19^101^ through the activation of glycogen synthase kinase 3β in prefrontal cortex and dorsal hippocampus^102^. Defects in formation of β-catenin causes disruption of bloodbrain barrier^103^ leading to the development of cerebrovascular thrombosis^104^, headache^105^, stroke^106^ and epileptic seizure^107^ during or aftermath of COVID-19. Stress-induced Dickkopf-1 protein formation prevents *CTNNB1* gene function in hippocampus thereby impairing memory^108^. Uncontrolled interactions of CTNNB1 with PSEN1^109^ and GLI2^110^ are linked to skin tumorigenesis, which may be suggestive for their possible involvement in COVID-19. Moreover, abnormalities in PSEN1 and GLI2 functions, associated with *CTNNB1* gene^111,112^ are likely to be implicated in developing Alzheimer’s disease- and holoprosencephaly-like features in COVID-19.

*ESR1* gene encodes estrogen receptor 1 that occurs primarily in the medial preoptic area and ventromedial nucleus of hypothalamus, which regulates diverse reproductive functions of both males and females^113^. ESR1 deems to share CTNNB1-^114^ and AKT1-^115^ mediated signaling pathways to accelerate cancer and neurodegeneration, respectively. Moreover, estrogen inhibits inflammation and immune responses in COVID-19 and reduces the COVID-19 susceptibility in females than in males, because of its higher concentration and greater number of ESR1 receptors in target tissues^116^.

In adult brain *PIK3CA* gene product PI3K via AKT1 pathway may exaggerate neurodegeneration in Alzheimer’s disease^117^, and FGFR1 dysregulation leads to ischemic stroke^118^ and holoprosencephaly^119^. Moreover, synchronized *PIK3CA* mutation and FGFR1 alteration are associated with *ESR1*-positive breast cancer^120^. Since COVID-19 develops inflammatory burst and lymphopenia, SARS-CoV-2 associated illness therefore may aggravate cancer prognosis^119^.

Notably, two prevalent genes *CTNNB1* and *AKT1* appeared to be common for both TN (Fig. 3a) and CGN (Fig. 4b). Both genes showed SSS-II (network specific semantic similarity score) values greater than threshold values in respective cases (Fig. 3e and Fig. 4e) and therefore functionally interlinked (Fig. 5f). *CTNNB1* appeared as the lone gene having both HB and diver node properties in TN. Interestingly, *CTNNB1* was the only gene which formed a ‘tripartite open network’ that linked with eight symptoms and those symptoms remained connected with eight diseases (Fig. 3a). *CTNNB1* in TN got connections with (a) five symptoms (viz. cerebral ischemia, vascular thrombosis, intracranial hypertension, seizures and epileptic seizures) in central nervous system (CNS), (b) two symptoms (viz. hypertonia and fatigue) in peripheral nervous system (PNS) and (c) one psychiatric symptom (viz. behavioral disorder). Moreover, it demonstrated that three symptoms connected with *CTNNB1* in the present tripartite network, also happened to occur in other diseases, co-infected with COVID-19, viz. (a) cerebral ischemia in alobar, lobar and semilobar holoprosencephaly, Behçet disease, early infantile epileptic encephalopathy, MELAS and meningioma; (b) vascular thrombosis in alobar, lobar and semilobar holoprosencephaly, amyotrophic lateral sclerosis and MELAS; (c) intracranial hypertension in MELAS. But no data is available yet about the rest of the five symptoms in any other diseases challenged none-ever with SARS-CoV-2. This suggests that certain neurological symptoms of COVID-19 are intermingled with other diseases and need special clinical attention.

## Acknowledgments

We thank Department of Biotechnology (DBT), Ministry of Science and Technology, Government of India, under BINC (Bioinformatics National Certification) scheme [No.: BT/BI/10/078/2014] for research fund. We also thank Professor Pritha Mukhopadhyay, coordinator of CPEPA-UGC centre [“Centre for Electrophysiological and Neuro-imaging studies including Mathematical Modelling” (CPEPA) through “University Grants Commission” (UGC)], under the University of Calcutta, India for providing necessary research support in centre. We thank Mr. Ritayan Chakrabarti for some mathematical comments.

## Author contributions

S. H., N. C., B. T., A. G. C. conceptualized, investigated, validated and resourced the study. The methodology was designed by S. H., B. T., N. C. All the Bioinformatic, statistical, formal and software analysis related to research data were performed by S. H. S.H. wrote the original draft and N. C., B. T., A. G. C. reviewed and edited the draft and all authors gave consent to publish the data presented in the article. The project study was administered by both corresponding authors N. C., B. T. and supervised by lead corresponding author N. C.

## Data and materials availability

All data are available in the paper.

## References and Notes

1. D. Wang, B. Hu, C. Hu, F. Zhu, X. Liu, J. Zhang, B. Wang, H. Xiang, Z. Cheng, Y. Xiong, Y. Zhao, Y. Li, X. Wang, Z. Peng, Clinical Characteristics of 138 Hospitalized Patients With 2019 Novel Coronavirus-Infected Pneumonia in Wuhan, China. JAMA, 2020, 323(11), 1061–1069, DOI: https://doi.org/10.1001/jama.2020.1585.

2. Y. Wang, Y. Wang, Y. Chen and Q. Qin, Unique epidemiological and clinical features of the emerging 2019 novel coronavirus pneumonia (COVID-19) implicate special control measures. J. Med. Virol., 2020, 92(6), 568–576, DOI: https://doi.org/10.1002/jmv.25748.

3. C. Huang, Y. Wang, X. Li, L. Ren, J. Zhao, Y. Hu, L. Zhang, G. Fan, J. Xu, X. Gu, Z. Cheng, T. Yu, J. Xia, Y. Wei, W. Wu, X. Xie, W. Yin, H. Li, M. Liu, Y. Xiao, H. Gao, L. Guo, J. Xie, G. Wang, R. Jiang, Z. Gao, Q. Jin, J. Wang and B. Cao, Clinical features of patients infected with 2019 novel coronavirus in Wuhan, China. Lancet, 2020, 395(10223), 497–506, DOI: https://doi.org/10.1016/S0140-6736(20)30183-5.

4. N. Zhu, D. Zhang, W. Wang, X. Li, B. Yang, J. Song, X. Zhao, B. Huang, W. Shi, R. Lu, P. Niu, F. Zhan, X. Ma, D. Wang, W. Xu, G. Wu, G. F. Gao and W. Tan, China Novel Coronavirus Investigating and Research Team, A Novel Coronavirus from Patients with Pneumonia in China, 2019. N. Engl. J. Med., 2020, 382(8), 727–733, DOI: https://doi.org/10.1056/NEJMoa2001017.

5. C. Wang, P. W. Horby, F. G. Hayden and G. F. Gao, A novel coronavirus outbreak of global health concern. Lancet, 2020, 395(10223), 470–473, DOI: https://doi.org/10.1016/S0140-6736(20)30185-9.

6. World Health Organization. Coronavirus disease 2019 (COVID-19): situation report, 2020, 94, DOI: https://www.who.int/docs/default-source/coronaviruse/situation-reports/20200423-sitrep-94-covid-19.pdf

7. M. Hoffmann, H. Kleine-Weber, S. Schroeder, N. Krüger, T. Herrler, S. Erichsen, T. S. Schiergens, G. Herrler, N. H. Wu, A. Nitsche, M. A. Müller, C. Drosten and S. Pöhlmann, SARS-CoV-2 Cell Entry Depends on ACE2 and TMPRSS2 and Is Blocked by a Clinically Proven Protease Inhibitor. Cell, 2020, 181(2), 271-280.e8, DOI: https://doi.org/10.1016/j.cell.2020.02.052.

8. V. J. Munster, M. Koopmans, N. van Doremalen, D. van Riel, E. de Wit, A Novel Coronavirus Emerging in China – Key Questions for Impact Assessment. N. Engl. J. Med., 2020, 382(8), 692–694, DOI: https://doi.org/10.1056/NEJMp2000929.

9. R. Chen, K. Wang, J. Yu, D. Howard, L. French, Z. Chen, C. Wen, Z. Xu, The Spatial and Cell-Type Distribution of SARS-CoV-2 Receptor ACE2 in the Human and Mouse Brains. Front. Neurol., 2021, 11, 573095, DOI: https://doi.org/10.3389/fneur.2020.573095.

10. S. Kremer, F. Lersy, M. Anheim, H. Merdji, M. Schenck, H. Oesterlé, F. Bolognini, J. Messie, A. Khalil, A. Gaudemer and S. Carré, Neurologic and neuroimaging findings in patients with COVID-19: A retrospective multicenter study. Neurology, 2020, 95(13), e1868–e1882, DOI: https://doi.org/10.1212/WNL.0000000000010112.

11. J. Wenzel, J. Lampe, H. Müller-Fielitz, R. Schuster, M. Zille, K. Müller, M. Krohn, J. Körbelin, L. Zhang, Ü. Özorhan and V. Neve, The SARS-CoV-2 main protease Mpro causes microvascular brain pathology by cleaving NEMO in brain endothelial cells. Nat. Neurosci., 2020, 24(11), 1522–1533, DOI: https://doi.org/10.1038/s41593-021-00926-1.

12. O. Al-Dalahmah, K. T. Thakur, A. S. Nordvig, M. L. Prust, W. Roth, A. Lignelli, A. C. Uhlemann, E. H. Miller, S. Kunnath-Velayudhan, A. Del Portillo and Y. Liu, Neuronophagia and microglial nodules in a SARS-CoV-2 patient with cerebellar hemorrhage. Acta Neuropathol. Commun., 2020, 8(1), 147, DOI: https://doi.org/10.1186/s40478-020-01024-2.

13. J. Meinhardt, J. Radke, C. Dittmayer, J. Franz, C. Thomas, R. Mothes, M. Laue, J. Schneider, S. Brünink, S. Greuel and M. Lehmann, Olfactory transmucosal SARS-CoV-2 invasion as a port of central nervous system entry in individuals with COVID-19. Nat. Neurosci., 2021, 24(2), 168–175, DOI: https://doi.org/10.1038/s41593-020-00758-5.

14. P. Guadarrama-Ortiz, J. A. Choreño-Parra, C. M. Sánchez-Martínez, F. J. Pacheco-Sánchez, A. I. Rodríguez-Nava and G. García-Quintero, Neurological Aspects of SARS-CoV-2 Infection: Mechanisms and Manifestations. Front. Neurol., 2020, 11, 1039, DOI: https://doi.org/10.3389/fneur.2020.01039.

15. M. Lima, V. Siokas, A. M. Aloizou, I. Liampas, A. F. A. Mentis, Z. Tsouris, A. Papadimitriou, P. D. Mitsias, A. Tsatsakis, D. P. Bogdanos and S. J. Baloyannis, Unraveling the Possible Routes of SARS-COV-2 Invasion into the Central Nervous System. Curr. Treat. Options Neurol., 2020, 22(11), 37, DOI: https://doi.org/10.1007/s11940-020-00647-z.

16. L. Pellegrini, A. Albecka, D. L. Mallery, M. J. Kellner, D. Paul, A. P. Carter, L. C. James and M. A. Lancaster, SARS-CoV-2 Infects the Brain Choroid Plexus and Disrupts the Blood-CSF Barrier in Human Brain Organoids. Cell stem cell, 2020, 27(6), 951-961.e5, DOI: https://doi.org/10.1016/j.stem.2020.10.001.

17. M. A. Ellul, L. Benjamin, B. Singh, S. Lant, B. D. Michael, A. Easton, R. Kneen, S. Defres, J. Sejvar and T. Solomon, Neurological associations of COVID-19. Lancet Neurol., 2020, 19(9), P767–P783, DOI: https://doi.org/10.1016/S1474-4422(20)30221-0.

18. T. Solomon, Neurological infection with SARS-CoV-2 – the story so far. Nat. Rev. Neurol., 2021, 17(2), 65–66, DOI: https://doi.org/10.1038/s41582-020-00453-w.

19. D. H. Brann, T. Tsukahara, C. Weinreb, M. Lipovsek, K. Van den Berge, B. Gong, R. Chance, I. C. Macaulay, H. J. Chou, R. B. Fletcher and D. Das, Non-neuronal expression of SARS-CoV-2 entry genes in the olfactory system suggests mechanisms underlying COVID-19-associated anosmia. Sci. Adv., 2020, 6(31), eabc5801, DOI: https://doi.org/10.1126/sciadv.abc5801.

20. M. E. Boroujeni, L. Simani, H. A. Bluyssen, H. R. Samadikhah, S. Zamanlui Benisi, S. Hassani, N. Akbari Dilmaghani, M. Fathi, K. Vakili, G. R. Mahmoudiasl and H. A. Abbaszadeh, Inflammatory Response Leads to Neuronal Death in Human Post-Mortem Cerebral Cortex in Patients with COVID-19. ACS Chem. Neurosci., 2021, 12(12), 2143–2150, DOI: https://doi.org/10.1021/acschemneuro.1c00111.

21. I. Mahalaxmi, J. Kaavya, S. Mohana Devi and V. Balachandar, COVID-19 and olfactory dysfunction: A possible associative approach towards neurodegenerative diseases. J. Cell. Physiol., 2021, 236(2), 763–770, DOI: https://doi.org/10.1002/jcp.29937.

22. M. A. Rahman, K. Islam, S. Rahman and M. Alamin, Neurobiochemical Cross-talk Between COVID-19 and Alzheimer’s Disease. Mol. Neurobiol., 2021, 58(3), 1017–1023, DOI: https://doi.org/10.1007/s12035-020-02177-w.

23. D. Sulzer, A. Antonini, V. Leta, A. Nordvig, R. J. Smeyne, J. E. Goldman, O. Al-Dalahmah, L. Zecca, A. Sette, L. Bubacco, O. Meucci, E. Moro, A. S. Harms, Y. Xu, S. Fahn and K. Ray Chaudhuri, COVID-19 and possible links with Parkinson’s disease and parkinsonism: from bench to bedside. NPJ Parkinsons Dis., 2020, 6, 18, DOI: https://doi.org/10.1038/s4153=-020-00123-0.

24. M. Bodro, Y. Compta and R. Sánchez-Valle, Presentations and mechanisms of CNS disorders related to COVID-19. Neurol. Neuroimmunol. Neuroinflamm., 2020, 8(1), e923, DOI: https://doi.org/10.1212/NXI.0000000000000923.

25. S. H. Y. Chou, E. Beghi, R. Helbok, E. Moro, J. Sampson, V. Altamirano, S. Mainali, C. Bassetti, J. I. Suarez, M. McNett and L. Nolan, Global Incidence of Neurological Manifestations Among Patients Hospitalized With COVID-19-A Report for the GCS-NeuroCOVID Consortium and the ENERGY Consortium. JAMA Netw. Open, 2021, 4(5), e2112131, DOI: https://doi.org/10.1001/jamanetworkopen.2021.12131.

26. J. P. Rogers, E. Chesney, D. Oliver, T. A. Pollak, P. McGuire, P. Fusar-Poli, M. S. Zandi, G. Lewis and A. S. David, Psychiatric and neuropsychiatric presentations associated with severe coronavirus infections: a systematic review and meta-analysis with comparison to the COVID-19 pandemic. Lancet Psychiat., 2020, 7(7), 611–627, DOI: https://doi.org/10.1016/S2215-0366(20)30203-0.

27. M. Taquet, J. R. Geddes, M. Husain, S. Luciano and P. J. Harrison, 6-month neurological and psychiatric outcomes in 236 379 survivors of COVID-19: a retrospective cohort study using electronic health records. Lancet Psychiat., 2021, 8(5), 416–427, DOI: https://doi.org/10.1016^2215-0366(21)00084-5.

28. R. Beyrouti, M. E. Adams, L. Benjamin, H. Cohen, S. F. Farmer, Y. Y. Goh, F. Humphries, H. R. Jäger, N. A. Losseff, R. J. Perry and S. Shah, Characteristics of ischaemic stroke associated with COVID-19. J. Neurol. Neurosurg. Psychiatry, 2020, 91(8), 889–891, DOI: http://doi.org/10.1136/jnnp-2020-323586.

29. T. Moriguchi, N. Harii, J. Goto, D. Harada, H. Sugawara, J. Takamino, M. Ueno, H. Sakata, K. Kondo, N. Myose and A. Nakao, A first case of meningitis/encephalitis associated with SARS-Coronavirus-2. Int. J. Infect. Dis., 2020, 94, 55–58, DOI: https://doi.org/10.1016/j.ijid.2020.03.062.

30. J. Helms, S. Kremer, H. Merdji, R. Clere-Jehl, M. Schenck, C. Kummerlen, O. Collange, C. Boulay, S. Fafi-Kremer, M. Ohana and M., Anheim, Neurologic Features in Severe SARS-CoV-2 Infection. N. Engl. J. Med., 2020, 382(23), 2268–2270, DOI: https://doi.org/10.1056/NEJMc2008597.

31. N. Poyiadji, G. Shahin, D. Noujaim, M. Stone, S. Patel and B. Griffith, COVID-19-associated acute hemorrhagic necrotizing encephalopathy: imaging features. Radiology, 2020, 296(2), E119–E120, DOI: https://doi.org/10.1148/radiol.2020201187.

32. R. W. Paterson, R. L. Brown, L. Benjamin, R. Nortley, S. Wiethoff, T. Bharucha, D. L. Jayaseelan, G. Kumar, R. E. Raftopoulos, L. Zambreanu and V. Vivekanandam, The emerging spectrum of COVID-19 neurology: clinical, radiological and laboratory findings. Brain, 2020, 143(10), 3104–3120, DOI: https://doi.org/10.1093/brain/awaa240.

33. S. Kremer, F. Lersy, M. Anheim, H. Merdji, M. Schenck, H. Oesterlé, F. Bolognini, J. Messie, A. Khalil, A. Gaudemer and S. Carré, Neurologic and neuroimaging findings in patients with COVID-19: A retrospective multicenter study. Neurology, 2020, 95(13), e1868–e1882, DOI: https://doi.org/10.1212/WNL.0000000000010112.

34. E. Pascual-Goñi, J. Fortea, A. Martínez-Domeño, N. Rabella, M. Tecame, C. Gómez-Oliva, L. Querol and B. Gómez-Ansón, COVID-19-associated ophthalmoparesis and hypothalamic involvement. Neurol. Neuroimmunol. Neuroinflamm., 2020, 7(5), e823, DOI: https://doi.org/10.1212/NXI.0000000000000823.

35. R. AlKetbi, D. AlNuaimi, M. AlMulla, N. AlTalai, M. Samir, N. Kumar and U. AlBastaki, Acute myelitis as a neurological complication of Covid-19: a case report and MRI findings. Radiol. Case Rep., 2020, 15(9), 1591–1595, DOI: https://doi.org/10.1016/j.radcr.2020.06.001.

36. J. Helms, S. Kremer, H. Merdji, M. Schenck, F. Severac, R. Clere-Jehl, A. Studer, M. Radosavljevic, C. Kummerlen, A. Monnier, C. Boulay, S. Fafi-Kremer, V. Castelain, M. Ohana, M. Anheim, F. Schneider and F. Meziani, Delirium and encephalopathy in severe COVID-19: a cohort analysis of ICU patients. Crit. care, 2020, 24(1), 491, DOI: https://doi.org/10.1186/s13054-020-03200-1.

37. T. H. Chua, Z. Xu and N. K. K. King, Neurological manifestations in COVID-19: a systematic review and meta-analysis. Brain Inj., 2020, 34(12), 1549–1568, DOI: https://doi.org/10.1080/02699052.2020.1831606.

38. F. Khatoon, K. Prasad and V. Kumar, Neurological manifestations of COVID-19: available evidences and a new paradigm. J Neurovirol., 2020, 26(5), 619–630, DOI: https://doi.org/10.1007/s13365-020-00895-4.

39. M. E. V. Collantes, Sy M. C. C. Espiritu AI, V. M. M. Anlacan and R. D. G. Jamora, Neurological Manifestations in COVID-19 Infection: A Systematic Review and Meta-Analysis. Can J Neurol Sci., 2021, 48(1), 66–76, DOI: https://doi.org/10.1017/cjn.2020.146.

40. T. T. Favas, P. Dev, R. N. Chaurasia, K. Chakravarty, R. Mishra, D. Joshi, V. N. Mishra, A., Kumar, V. K. Singh, M. Pandey and A. Pathak, Neurological manifestations of COVID-19: a systematic review and meta-analysis of proportions. Neurol Sci., 2020, 41(12), 3437–3470, DOI: https://doi.org/10.1007/s10072-020-04801-y.

41. G. Nepal, J. H. Rehrig, G. S. Shrestha, Y. K. Shing, J. K. Yadav, R. Ojha, G. Pokhrel, Z. L. Tu and D. Y. Huang, Neurological manifestations of COVID-19: a systematic review. Crit. Care, 2020, 24, 421, DOI: https://doi.org/10.1186/s13054-020-03121-z.

42. D. Vitalakumar, A. Sharma, A. Kumar and S. J. S. Flora, Neurological Manifestations in COVID-19 Patients: A Meta-Analysis. ACS Chem. Neurosci., 2021, 12(15), 2776–2797, DOI: https://doi.org/10.1021/acschemneuro.1c00353.

43. S. Bagnato, C. Boccagni, G. Marino, C. Prestandrea, T. D’Agostino and F. Rubino, Critical illness myopathy after COVID-19. Int. J. Infect. Dis., 2020, 99, 276–278, DOI: https://doi.org/10.1016/j.ijid.2020.07.072.

44. L. Mao, H. Jin, M. Wang, Y. Hu, S. Chen, Q. He, J. Chang, C. Hong, Y. Zhou, D. Wang and X. Miao, Neurologic Manifestations of Hospitalized Patients With Coronavirus Disease 2019 in Wuhan, China. JAMA Neurol., 2020, 77(6), 683–690, DOI: https://doi.org/10.1001/jamaneurol.2020.1127.

45. H. Zhao, D. Shen, H. Zhou, J. Liu and S. Chen, Guillain-Barré syndrome associated with SARS-CoV-2 infection: causality or coincidence?. Lancet Neurol., 2020, 19(5), 383–384, DOI: https://doi.org/10.1016/S1474-4422(20)30109-5.

46. K. Prasad, S. Y. AlOmar, S. A. M. Alqahtani, M. Z. Malik and V. Kumar, Brain Disease Network Analysis to Elucidate the Neurological Manifestations of COVID-19. Mol. Neurobiol., 2021, 58(5), 1875–1893, DOI: https://doi.org/10.1007/s12035-020-02266-w.

47. Q. Wu, X. Coumoul, P. Grandjean, R. Barouki and K. Audouze, Endocrine disrupting chemicals and COVID-19 relationships: A computational systems biology approach. Environ. Int., 2020, 157, 106232, DOI: https://doi.org/10.1016/j.envint.2020.106232.

48. A. Halu, M. De Domenico, A. Arenas and A. Sharma, The multiplex network of human diseases. NPJ Syst. Biol. Appl., 2019, 5, 15, DOI: https://doi.org/10.1038/s41540-019-0092-5.

49. A. Sepehrinezhad, F. Rezaeitalab, A. Shahbazi and S. Sahab-Negah, A Computational-Based Drug Repurposing Method Targeting SARS-CoV-2 and its Neurological Manifestations Genes and Signaling Pathways. Bioinform. Biol. Insights., 2021, 15, 11779322211026728, DOI: https://doi.org/10.1177/11779322211026728.

50. D. Moher, A. Liberati, J. Tetzlaff and D. G. Altman, PRISMA Group Preferred reporting items for systematic reviews and meta-analysis: the PRISMA statement. Int. J. Surg., 2010, 8(5), 336–341, DOI: https://doi.org/10.1371/journal.pmed.1000097.

51. S. Köhler, M. Gargano, N. Matentzoglu, L. C. Carmody, D. Lewis-Smith, N. A. Vasilevsky, D. Danis, G. Balagura, G. Baynam, A. M. Brower, T. J. Callahan, C. G. Chute, J. L. Est, P. D. Galer, S. Ganesan, M. Griese, M. Haimel, J. Pazmandi, M. Hanauer, N. L. Harris, M. J. Hartnett, M. Hastreiter, F. Hauck, Y. He, T. Jeske, H. Kearney, G. Kindle, C. Klein, K. Knoflach, R. Krause, D. Lagorce, J. A. McMurry, J. A. Miller, M. C. Munoz-Torres, R. L. Peters, C. K. Rapp, A. M. Rath, S. A. Rind, A. Z. Rosenberg, M. M. Segal, M. G. Seidel, D. Smedley, T. Talmy, Y. Thomas, S. A. Wiafe, J., Xian Z. Yüksel, I. Helbig, C. J. Mungall, M. A. Haendel and P. N. Robinson, The Human Phenotype Ontology in 2021, Nucleic Acids Res., 2021, 49(D1), D1207–D1217, DOI: https://doi.org/10.1093/nar/gkaa1043.

52. B. Pesta, J. Fuerst and E. O. W. Kirkegaard, Bibliometric Keyword Analysis across Seventeen Years (2000-2016) of Intelligence Articles. J Intell., 2018, 6(4), 46, DOI: https://doi.org/10.3390/jintelligence6040046.

53. L. Deng, D. Ye, J. Zhao and J. Zhang, MultiSourcDSim: an integrated approach for exploring disease similarity. BMC Medical Inform. Decis. Mak., 2019, 19(Suppl 6), 269, DOI: https://doi.org/10.1186/s12911-019-0968-8.

54. N. Rappaport, M. Twik, I. Plaschkes, R. Nudel, T. Iny Stein, J. Levitt, M. Gershoni, C. P. Morrey, M. Safran and D. Lancet, MalaCards: an amalgamated human disease compendium with diverse clinical and genetic annotation and structured search. Nucleic Acids Res., 2017, 45(D1), D877–D887, DOI: https://doi.org/10.1093/nar/gkw1012.

55. X. Zhou, J. Menche, A. L. Barabási and A. Sharma, Human symptoms-disease network. Nat. Commun., 2014, 5, 4212, DOI: https://doi.org/10.1038/ncomms5212.

56. D. Szklarczyk, A. L. Gable, K. C. Nastou, D. Lyon, R. Kirsch, S. Pyysalo, N. T. Doncheva, M. Legeay, T. Fang, P. Bork and L. J. Jensen, The STRING database in 2021: customizable protein-protein networks, and functional characterization of user-uploaded gene/measurement sets. Nucleic Acids Res., 2021, 49(D1), D605–D612, DOI: https://doi.org/10.1093/nar/gkaa1074.

57. P. Shannon, A. Markiel, O. Ozier, N. S. Baliga, J. T. Wang, D. Ramage, N. Amin, B. Schwikowski, T. Ideker, Cytoscape: a software environment for integrated models of biomolecular interaction networks. Genome Res., 2003, 13(11), 2498–2504, DOI: https://doi.org/10.1101/gr.1239303.

58. G. Scardoni, M. Petterlini and C. Laudanna, Analyzing biological network parameters with CentiScaPe. Bioinformatics, 2009, 25(21), 2857–2859, DOI: https://doi.org/10.1093/bioinformatics/btp517.

59. S. Gagliardi, T. E. Poloni, C. Pandini, M. Garofalo, F. Dragoni, V. Medici, A. Davin, S. D. Visonà, M. Moretti, D. Sproviero and O. Pansarasa, Detection of SARS-CoV-2 genome and whole transcriptome sequencing in frontal cortex of COVID-19 patients. Brain Behav. Immun., 2021, 97, 13–21, DOI: https://doi.org/10.1016/j.bbi.2021.05.012.

60. A. B. Owen, J. Stuart, K. Mach, A. M. Villeneuve and S. Kim, A gene recommender algorithm to identify coexpressed genes in C. elegans. Genome Res., 2003, 13(8), 1828–1837, DOI: https://doi.org/10.1101/gr.1125403.

61. G. J. Hather, A. B. Owen and T. P. Speed, geneRecommender: A gene recommender algorithm to identify genes coexpressed with a query set of genes. R package v1.64.0. (2021), DOI: https://bioconductor.org/packages/3.14/bioc/html/geneRecommender.html.

62. P. E. Meyer, F. Lafitte and G. Bontempi, minet: A R/Bioconductor Package for Inferring Large Transcriptional Networks Using Mutual Information. BMC Bioinformatics, 2008, 9, 461, DOI: https://doi.org/10.1186/1471-2105-9-461.

63. A. A. Margolin, I. Nemenman, K. Basso, C. Wiggins, G. Stolovitzky, R. Dalla Favera, A. Califano, ARACNE: an algorithm for the reconstruction of gene regulatory networks in a mammalian cellular context. BMC Bioinform., 2006, 7(Suppl 1), S7, DOI: https://doi.org/10.1186/1471-2105-7-S1-S7.

64. J. D. J. Han, N. Bertin, T. Hao, D. S. Goldberg, G .F. Berriz, L. V. Zhang, D. Dupuy, A. J. Walhout, M. E. Cusick, F. P. Roth and M. Vidal, Evidence for dynamically organized modularity in the yeast protein-protein interaction network. Nature, 2004, 430(6995), 88–93, DOI: https://doi.org/10.1038/nature02555.

65. H. Yu, P. Braun, M. A. Yildirim, I. Lemmens, K. Venkatesan, J. Sahalie, T. Hirozane-Kishikawa, F. Gebreab, N. Li, N. Simonis, and T. Hao, High-quality binary protein interaction map of the yeast interactome network. Science, 2008, 322(5898), 104–110, DOI: https://doi.org/10.1126/science.1158684.

66. S. Agarwal, C. M. Deane, M. A. Porter, N. S. Jones, Revisiting date and party hubs: novel approaches to role assignment in protein interaction networks. PLoS Comput. Biol., 2010, 6(6), e1000817, DOI: https://doi.org/10.1371/journal.pcbi.1000817.

67. X., Chang, T. Xu, Y. Li and K. Wang, Dynamic modular architecture of protein-protein interaction networks beyond the dichotomy of ‘date’ and ‘party’ hubs. Sci. Rep., 2013, 3, 1691, DOI: https://doi.org/10.1038/srep01691.

68. L. Wu, M. Li, J. Wang and F. X. Wu, CytoCtrlAnalyser: a Cytoscape app for biomolecular network controllability analysis. Bioinformatics, 2018, 34(8), 1428–1430, DOI: https://doi.org/10.1093/bioinformatics/btx764.

69. A. Vinayagam, T. E. Gibson, H. J. Lee, B. Yilmazel, C. Roesel, Y. Hu, Y. Kwon, A. Sharma, Y. Y. Liu, N. Perrimon and A. L. Barabási, Controllability analysis of the directed human protein interaction network identifies disease genes and drug targets. Proc. Natl. Acad. Sci. U.S.A., 2016, 113(18), 4976–4981, DOI: https://doi.org/10.1073/pnas.1603992113.

70. Y. Y. Liu, J. J. Slotine and A. L. Barabási, Controllability of complex networks. Nature, 2011, 473, 167–173, DOI: https://doi.org/10.1038/nature10011.

71. V. Ravindran, J. C. Nacher, T. Akutsu, M. Ishitsuka, A. Osadcenco, V. Sunitha, G. Bagler, J. M. Schwartz and D. L. Robertson, Network controllability analysis of intracellular signalling reveals viruses are actively controlling molecular systems. Sci. Rep., 2019, 9(1), 2066, DOI: https://doi.org/10.1038/s41598-018-38224-9.

72. J. Loscalzo and A. L. Barabási, Systems biology and the future of medicine. Wiley Interdiscip. Rev. Syst. Biol. Med., 2011, 3(6), 619–627, DOI: https://doi.org/10.1002%2Fwsbm.144.

73. J. B. Bard and S. Y. Rhee, Ontologies in biology: design, applications and future challenges. Nat. Rev. Genet., 2004, 5(3), 213–222, DOI: https://doi.org/10.1038/nrg1295.

74. S. Y. Rhee, V. Wood, K. Dolinski and S. Draghici, Use and misuse of the gene ontology annotations. Nat. Rev. Genet., 2008, 9(7), 509–515, DOI: https://doi.org/10.1038/nrg2363.

75. G. Yu, F. Li, Y. Qin, X. Bo, Y. Wu and S. Wang, GOSemSim: an R package for measuring semantic similarity among GO terms and gene products. Bioinformatics, 2010, 26(7), 976–978, DOI: https://doi.org/10.1093/bioinformatics/btq064.

76. J. Z. Wang, Z. Du, R. Payattakool, P. S. Yu and C. F. Chen, A new method to measure the semantic similarity of GO terms. Bioinformatics, 2007, 23(10), 1274–1281, DOI: https://doi.org/10.1093/bioinformatics/btm087.

77. C. Pesquita, D. Faria, A. O. Falcão, P. Lord and F. M. Couto, Semantic similarity in biomedical ontologies. PLoS Comput. Biol., 2009, 5(7), e1000443, DOI: https://doi.org/10.1371/journal.pcbi.1000443.

78. X. Robin, N. Turck, A. Hainard, N. Tiberti, F. Lisacek, J. C. Sanchez and M. Müller, pROC: an open-source package for R and S+ to analyze and compare ROC curves. BMC Bioinform., 2011, 12, 77, DOI: https://doi.org/10.1186/1471-2105-12-77.

79. E. Y. Chen, C. M. Tan, Y. Kou, Q. Duan, Z. Wang, G. V. Meirelles, N. R. Clark, A. Ma’ayan, Enrichr: interactive and collaborative HTML5 gene list enrichment analysis tool. BMC Bioinform., 2013, 14, 128, DOI: https://doi.org/10.1186/1471-2105-14-128.

80. Z. Xie, A. Bailey, M. V. Kuleshov, D. J. Clarke, J. E. Evangelista, S. L. Jenkins, A. Lachmann, M. L. Wojciechowicz, E. Kropiwnicki, K. M. Jagodnik and M. Jeon, Gene Set Knowledge Discovery with Enrichr. Curr. Protoc., 2021, 1(3), e90, DOI: https://doi.org/10.1002/cpz1.90.

81. P. C. Fernando, P. M. Mabee and E. Zeng, Integration of anatomy ontology data with protein-protein interaction networks improves the candidate gene prediction accuracy for anatomical entities. BMC Bioinform., 2020, 21(1), 442, DOI: https://doi.org/10.1186/s12859-020-03773-2.

82. P. Devkota, M. C. Danzi and S. Wuchty, Beyond degree and betweenness centrality: Alternative topological measures to predict viral targets. PloS One, 2018, 13(5), e0197595, DOI: https://doi.org/10.1371/journal.pone.0197595.

83. GO-Consortium. The Gene Ontology (GO) database and informatics resource. Nucleic Acids Res., 2004, 32(Database issue), D258–D261, DOI: https://doi.org/10.1093/nar/gkh036.

84. J. Peng, H. Wang, J. Lu, W. Hui, Y. Wang and X. Shang, Identifying term relations cross different gene ontology categories. BMC Bioinform., 2017, 18(Suppl 16), 573, DOI: https://doi.org/10.1186/s12859-017-1959-3.

85. W. Y. Hwang, Biological feature selection and disease gene identification using new stepwise random forests. Ind. Eng. Manag. Syst., 2017, 16(1), 64–79, DOI: https://doi.org/10.7232/iems.2017.16.1.064.

86. S. Chavali, F. Barrenas, K. Kanduri and M. Benson, Network properties of human disease genes with pleiotropic effects. BMC Syst. Biol., 2010, 4, 78, DOI: https://doi.org/10.1186/1752-0509-4-78.

87. K. I. Goh, M. E. Cusick, D. Valle, B. Childs, M. Vidal and A. L. Barabási, The human disease network. Proc. Natl. Acad. Sci. U.S.A., 2007, 104(21), 8685–8690, DOI: https://doi.org/10.1073pnas.0701361104.

88. K. Lage, N. T. Hansen, E. O. Karlberg, A. C. Eklund, F. S. Roque, P. K. Donahoe, Z. Szallasi, T. S. Jensen and S. Brunak, A large-scale analysis of tissue-specific pathology and gene expression of human disease genes and complexes. Proc. Natl. Acad. Sci. U.S.A., 2008, 105(52), 20870–20875, DOI: https://doi.org/10.1073pnas.0810772105.

89. S. van Dam, U. Võsa, A. van der Graaf, L. Franke and J. P. de Magalhães, Gene coexpression analysis for functional classification and gene-disease predictions. Brief. Bioinform., 2018, 19(4), 575–592, DOI: https://doi.org/10.1093/bib/bbw139.

90. J. Wang, M. Li, Y. Deng and Y. Pan, Recent advances in clustering methods for protein interaction networks. BMC Genom., 2010, 11(Suppl 3), 1–19, DOI: https://doi.org/10.1186/1471-2164-11-S3-S10.

91. B. Schreiber, A. Patel and A. Verma, Shedding light on COVID-19: ADAM17 the missing link? Am. J. Ther, 2021, 28(3), e358–e360, DOI: https://doi.org/10.1097/MJT.0000000000001226.

92. M. Qian, X. Shen and H. Wang, The Distinct Role of ADAM17 in APP Proteolysis and Microglial Activation Related to Alzheimer’s Disease. Cell. Mol. Neurobiol., 2016, 36, 471–482, DOI: https://doi.org/10.1007/s10571-015-0232-4.

93. H. Wang, S. Ma, J. Li, M. Zhao, X. Huo, J. Sun, L. Sun, J. Hu and Q. Liu, ADAM17 participates in the protective effect of paeoniflorin on mouse brain microvascular endothelial cells. J. Cell. Physiol., 2018, 233(12), 9320–9329, DOI: https://doi.org/10.1002/jcp.26308.

94. I. Bernard, D. Limonta, L. K. Mahal and T. C. Hobman, Endothelium Infection and Dysregulation by SARS-CoV-2: Evidence and Caveats in COVID-19. Viruses, 2020, 13(1), 29, DOI: https://doi.org/10.3390/v13010029.

95. A. B. Dixit, A. Srivastava, D. Sharma, M. Tripathi, D. Paul, S. Lalwani, R. Doddamani, M. C. Sharma, J. Banerjee and P. S. Chandra, Integrated Genome-Wide DNA Methylation and RNAseq Analysis of Hippocampal Specimens Identifies Potential Candidate Genes and Aberrant Signalling Pathways in Patients with Hippocampal Sclerosis. Neurol. India, 2020, 68(2), 307–313, DOI: https://doi.org/10.4103/0028-3886.280649.

96. G. Stepanova, Biologia Futura: is ADAM 17 the reason for COVID-19 susceptibility in hyperglycemic and diabetic patients?. Biol. Futura, 2021, 72(3), 291–297, DOI: https://doi.org/10.1007/s42977-021-00092-2.

97. H. J. Lim, P. Crowe and J. L. Yang, Current clinical regulation of PI3K/PTEN/Akt/mTOR signalling in treatment of human cancer. J Cancer Res. Clin. Oncol., 2015, 141(4), 671–689, DOI: https://doi.org/10.1007/s00432-014-1803-3.

98. H. H. Wong, P. Kumar, F. P. L. Tay, D. Moreau, D. X. Liu and F. Bard, Genome-Wide Screen Reveals Valosin-Containing Protein Requirement for Coronavirus Exit from Endosomes. J. Virol, 2015, 89(21), 11116–11128, DOI: https://doi.org/10.1128%2FJVI.01360-15.

99. Y. Zhou, J. Xu, Y. Hou, J. B. Leverenz, A. Kallianpur, R. Mehra, Y. Liu, H. Yu, A. A. Pieper, L. Jehi and F. Cheng, Network medicine links SARS-CoV-2/COVID-19 infection to brain microvascular injury and neuroinflammation in dementia-like cognitive impairment. Alzheimers Res. Ther., 2021, 13(1), 110, DOI: https://doi.org/10.1186/s13195-021-00850-3.

100. L. Pizzamiglio, E. Focchi, C. Cambria, L. Ponzoni, S. Ferrara, F. Bifari, G. Desiato, N. Landsberger, L. Murru, M. Passafaro and M. Sala, The DNA repair protein ATM as a target in autism spectrum disorder. JCI Insight, 2021, 6(3), e133654, DOI: https://doi.org/10.1172%2Fjci.insight.133654.

101. B. Vastrad, C. Vastrad and A. Tengli, Bioinformatics analyses of significant genes, related pathways, and candidate diagnostic biomarkers and molecular targets in SARS-CoV-2/COVID-19. Gene Rep., 2020, 21, 100956, DOI: https://doi.org/10.1016%2Fj.genrep.2020.100956.

102. L. Z. Xu, D. F. Xu, Y. Han, L. J. Liu, C. Y. Sun, J. H. Deng, R. X. Zhang, M. Yuan, S. Z. Zhang, Z. M. Li and Y. Xu, BDNF-GSK-3β-β-Catenin Pathway in the mPFC Is Involved in Antidepressant-Like Effects of Morinda officinalis Oligosaccharides in Rats. Int. J. Neuropsychopharmacol., 2017, 20(1), 83–93, DOI: https://doi.org/10.1093/ijnp/pyw088.

103. J. H. Fosse, G. Haraldsen, K. Falk and R. Edelmann, Endothelial Cells in Emerging Viral Infections. Front. Cardiovasc. Med., 2021, 8, 619690, DOI: https://doi.org/10.3389/fcvm.2021.619690.

104. Y. Fu, Y. Cheng and Y. Wu, Understanding SARS-CoV-2-Mediated Inflammatory Responses: From Mechanisms to Potential Therapeutic Tools. Virol. Sin., 2020, 35(3), 266–271, DOI: https://doi.org/10.1007/s12250-020-00207-4.

105. Y. Wu, X. Xu, Z. Chen, J. Duan, K. Hashimoto, L. Yang, C. Liu and C. Yang, Nervous system involvement after infection with COVID-19 and other coronaviruses. Brain Behav. Immun., 2020, 87, 18–22, DOI: https://doi.org/10.1016%2F3j.bbi.2020.03.031.

106. I. Cappuccio, A. Calderone, C. L. Busceti, F. Biagioni, F. Pontarelli, V. Bruno, M. Storto, G. T. Terstappen, G. Gaviraghi, F. Fornai and G. Battaglia, Induction of Dickkopf-1, a negative modulator of the Wnt pathway, is required for the development of ischemic neuronal death. J. Neurosci., 2005, 25(10), 2647–2657, DOI: https://doi.org/10.1523/JNEUROSCI.5230-04.2005.

107. J. Theilhaber, S. N. Rakhade, J. Sudhalter, N. Kothari, P. Klein, J. Pollard and F. E. Jensen, Gene expression profiling of a hypoxic seizure model of epilepsy suggests a role for mTOR and Wnt signaling in epileptogenesis. PloS One, 2013, 8(9), e74428. DOI: https://doi.org/10.1371/journal.pone.0074428.

108. F. Matrisciano, C. L. Busceti, D. Bucci, R. Orlando, A. Caruso, G. Molinaro, I. Cappuccio, B. Riozzi, R. Gradini, M. Motolese and F. Caraci, Induction of the Wnt antagonist Dickkopf-1 is involved in stress-induced hippocampal damage. PloS One, 2011, 6(1), e16447, DOI: https://doi.org/10.1371/journal.pone.0016447.

109. X. Xia, S. Qian, S. Soriano, Y. Wu, A. M. Fletcher, X. J. Wang, E. H. Koo, X. Wu and H. Zheng, Loss of presenilin 1 is associated with enhanced beta-catenin signaling and skin tumorigenesis. Proc. Natl. Acad. Sci. U.S.A., 2001, 98(19), 10863–10868, DOI: https://doi.org/10.1073/pnas.191284198.

110. E. Pantazi, E. Gemenetzidis, M. T. Teh, S. V. Reddy, G. Warnes, C. Evagora, G. Trigiante and M. P. Philpott, GLI2 Is a Regulator of β-Catenin and Is Associated with Loss of E-Cadherin, Cell Invasiveness, and Long-Term Epidermal Regeneration. J. Invest. Dermatol., 2017, 137(8), 1719–1730, DOI: https://doi.org/10.1016/j.jid.2016.11.046.

111. E. Roessler, Y. Z. Du, J. L. Mullor, E. Casas, W. P. Allen, G. Gillessen-Kaesbach, E. R. Roeder, J. E. Ming, A. R. i Altaba and M. Muenke, Loss-of-function mutations in the human GLI2 gene are associated with pituitary anomalies and holoprosencephaly-like features. Proc. Natl. Acad. Sci. U.S.A., 2003, 100(23), 13424–13429, DOI: https://doi.org/10.1073pnas.2235734100.

112. A. M. Dahlin, M. V. Hollegaard, C. Wibom, U. Andersson, D. M. Hougaard, I. Deltour, U. Hjalmars and B. Melin, CCND2, CTNNB1, DDX3X, GLI2, SMARCA4, MYC, MYCN, PTCH1, TP53, and MLL2 gene variants and risk of childhood medulloblastoma. J. Neurooncol., 2015, 125(1), 75–78, DOI: https://doi.org/10.1007/s11060-015-1891-1.

113. M. B. Yilmaz, A. Wolfe, Y. H. Cheng, C. Glidewell-Kenney, J. L. Jameson and S. E. Bulun, Aromatase promoter I.f is regulated by estrogen receptor alpha (ESR1) in mouse hypothalamic neuronal cell lines. Biol. Reprod., 2009, 81(5), 956–965, DOI: https://doi.org/10.1095/biolreprod.109.077206.

114. D. Barh, M. E. García-Solano, S. Tiwari, A. Bhattacharya, N. Jain, D. Torres-Moreno, B. Ferri, A. Silva, V. Azevedo, P. Ghosh and K. Blum, BARHL1 Is Downregulated in Alzheimer’s Disease and May Regulate Cognitive Functions through ESR1 and Multiple Pathways. Genes, 2017, 8(10), 245, DOI: https://doi.org/10.3390/genes8100245.

115. A. S. Khatpe, A. K. Adebayo, C. A. Herodotou, B. Kumar and H. Nakshatri, Nexus between PI3K/AKT and Estrogen Receptor Signaling in Breast Cancer. Cancers, 2021, 13(3), 369, DOI: https://doi.org/10.3390/cancers13030369.

116. F. Li, A. C. Boon, A. P. Michelson, R. E. Foraker, M. Zhan and P. R. Payne, Estrogen Hormone Is an Essential Sex Factor Inhibiting Inflammation and Immune Response in COVID-19. 2021, Res. Sq., rs.3.rs–936900, DOI: https://doi.org/10.21203/rs.3.rs-936900/v1.

117. S. C. Hopp, Y. Lin, D. Oakley, A. D. Roe, S. L. DeVos, D. Hanlon and B. T. Hyman, The role of microglia in processing and spreading of bioactive tau seeds in Alzheimer’s disease. J. Neuroinflammation, 2018, 15(1), 269, DOI: https://doi.org/10.1186/s12974-018-1309-z.

118. D. Wang, F. Liu, L. Zhu, P. Lin, F. Han, X. Wang, X. Tan, L. Lin and Y. Xiong, FGF21 alleviates neuroinflammation following ischemic stroke by modulating the temporal and spatial dynamics of microglia/macrophages. J. Neuroinflammation, 2020, 17(1), 257, DOI: https://doi.org/10.1186/s12974-020-01921-2.

119. C. Dubourg, W. Carré, H. Hamdi-Rozé, C. Mouden, J. Roume, B. Abdelmajid, D. Amram, C. Baumann, N. Chassaing, C. Coubes and L. Faivre-Olivier, Mutational Spectrum in Holoprosencephaly Shows That FGF is a New Major Signaling Pathway. Hum. Mutat., 2016, 37(12), 1329–1339, DOI: https://doi.org/10.1002/humu.23038.

120. D. M. Hyman, B. Tran, L. Paz-Ares, J. P. Machiels, J. H. Schellens, P. L. Bedard, M. Campone, P. A. Cassier, J. Sarantopoulos, U. Vaishampayan and R. Chugh, Combined PIK3CA and FGFR inhibition with alpelisib and infigratinib in patients with PIK3CA-mutant solid tumors, with or without FGFR alterations. JCO Precis. Oncol., 2019, 3, 1–13, DOI: http://ascopubs.org/doi/full/10.1200/PO.19.00221.

